# Integrating metagenome-scale metabolic modelling and metabolomics to identify biochemical interactions in *Microcystis* phycospheres

**DOI:** 10.64898/2026.03.18.712574

**Authors:** Juliette Audemard, Nicolas Creusot, Julie Leloup, Charlotte Duval, Sébastien Halary, Lou Mary, Melissa Eon, Thomas Forjonel, Mohamed Mouffok, Rémy Puppo, Elodie Belmonte, Veronique Gautier, Jeanne Got, Marie Lefebvre, Gabriel V. Markov, Coralie Muller, Benjamin Marie, Binta Diémé, Clémence Frioux

## Abstract

Favoured by global changes, freshwater cyanobacterial harmful blooms generate major ecological, economical and public health challenges. *Microcystis*, one of the most widespread cyanobacterial genera, grows within a phycosphere where specialised interactions with its microbiome occur, and are suspected to influence bloom appearance and its potential toxicity. Using a combination of metagenomic, metabolomic and metabolic modelling, we characterised the phycospheres of twelve *Microcystis* strains isolated from a French pond. The distribution of metabolic reactions within *Microcystis* was consistent with their genospecies, whereas the metabolic landscape at the community level diverged from cyanobacterial phylogeny indicating functional decoupling between cyanobacteria and their associated microbiomes. Phycosphere-associated bacteria substantially expand the metabolic repertoire of the system, while maintaining functional redundancy within and across communities. On the other hand, metabolomic profiles were largely driven by cyanobacterial metabolic outputs. Metabolic modelling, together with the identification of toxic specialised metabolites produced by specific biosynthetic gene clusters, further highlighted differences in metabolic potential among phycospheres. Together, these findings deepen the understanding of *Microcystis*’ phycosphere functioning, demonstrate the value of multi-omics systems biology approaches, and underscore the ecological relevance of interspecies and inter-phycosphere metabolic interactions as a structuring process in bloom-associated microbiomes.

## 1 Introduction

Freshwater harmful cyanobacterial blooms (HCBs) have become a globally recurring phenomenon, mainly driven by rising temperatures and increased nutrient loads from anthropogenic activities [65]. These blooms, characterised by the proliferation of cyanobacteria such as *Microcystis*, pose major ecological and societal challenges through the release of large amounts of organic matter and toxic metabolites (*e.g.* microcystins). Such inputs disrupt ecosystem functioning with severe consequences for biodiversity, public health, and local economies [42, 70]. Despite substantial progress in understanding the environmental drivers and the dynamics of cyanobacterial blooms, these phenomena are still difficult to predict and overcome.

A particularly understudied aspect of these phenomena is the complex network of metabolic interactions within the *phycosphere* - the microscale environment surrounding phytoplankton cells and aggregates - colonised by heterotrophic bacteria. While the taxonomic composition of phycosphere-associated microbiomes varies with cyanobacterial species [56], their global metabolic potential appears conserved, suggesting functional redundancy [68]. These microbial interactions within the phycosphere are thought to support phytoplankton growth, stress tolerance and bloom persistence through nutrient recycling and metabolite exchange, involving commensalism and competition [44, 55, 10, 82]. Yet, precise mechanisms and consequences of this microbial syntrophy remain poorly understood [22].

Addressing these knowledge gaps, a promising approach is to study simplified, cultivated, or synthetic microbial communities [18], as non-axenic cultures of phytoplankton host a specific but reduced microbiome [88]. In this context, genome-scale metabolic networks (GSMNs) are levers to generate hypotheses about functional roles within microbial communities, and connect genes, proteins, and biochemical reactions [14, 4]. However, applications to freshwater cyanobacterial communities remain limited, with most studies focusing on marine systems or isolated cyanobacteria [69, 86, 72]. Metabolomics complements these approaches by detecting main metabolites potentially mediating microbial interactions and refining metabolic model predictions [20, 26, 23, 74, 64].

Here, we present a systems biology-based characterisation of the taxonomic composition and co-metabolism of 12 cultivated phycospheres of *Microcystis* spp., isolated from a eutrophic freshwater pond during a bloom event. By integrating metagenomics, genome-scale metabolic modelling and metabolomics, we investigate the metabolic capabilities of *Microcystis* strains and their associated microbiomes. This combined approach provides a functional view of the phycosphere organisation, and reveals putative metabolic interactions and community-specific functions.

## 2 Material and Methods

### 2.1 Phycosphere isolation and culture

Water samples were collected in September 2021, from a suburban freshwater pond near Paris, Ilede-France (Cergy-Pontoise, GPS coordinates lat. 49.01343 long. 2.02575), experiencing a *Microcystis* bloom, (chlorophyl-content ≈ 10 *µ*g.L*^−^*^1^)[34].

After pre-filtration on 50 µm porosity, twelve monoclonal *Microcystis* strains and their associated bacterial consortia were isolated by repeated of single-cell or small-colony transfers on solid or liquid media under an inverted microscope (NIKON ECLIPSE TS100, Japan). Viable clones were then cultivated in 25 cm^2^ culture flasks (CORNING-FALCON) containing 10 mL of Z8 medium. Cyanobacterial strains were maintained in the Paris Museum Collection (PMC, https://mcam.mnhn.fr/en/ cyanobacteria-and-live-microalgae-470) at 18*^◦^*C, using white LED-powered lights providing an irradiance of 8-10 µm photons/m^2^/s, with a photoperiod of 13h light/11h dark. Isolated strains and cultures were all monoclonal and non-axenic. They were then cultivated in BG11 medium, with a photoperiod of 16h light/8h dark and ∼14 µm photons/m^2^/s. After optimal growth, cultures were centrifuged to separately collect cell pellets and supernatants for genomic and metabolomic analyses. Two complementary extraction protocols were used for metabolomic analyses to maximize coverage (see section 2.8).

### 2.2 Shotgun metagenomic sequencing

DNA was extracted from lyophilised cultures (50 mg) [30] using the DNeasy PowerLyzer PowerSoil kit (Qiagen) following an adapted protocol (manufacturer’s instructions from step 5) without bead beating [76]. Cells were lysed with PowerBead Solution (750 µL) and Solution C1 (60 µL) by two 5-minutes incubation at 70°C with 30s of vortexing before and after each incubation. Long-read shotgun metagenomic sequencing was performed using a PacBio Revio platform (Pacific Biosciences, Menlo Park, CA, USA) at the Gentyane sequencing facility (Clermont-Ferrand, France) with detailed protocols in Supplementary Material and Methods.

### 2.3 Metagenomic assembly and binning

Reads were assembled using metaMDBG (v1.1) [5], and binned with metaBAT 2 (v2.12.1) [46]. Bin quality was assessed with CheckM2 [17] (v1.1.0). High quality (HQ) metagenome-assembled genomes (MAGs, completeness *>*90%, contamination *<*5%) were taxonomically assigned using GTDB-tk (v2.4.1) [16] with the r226 GTDB release as reference [66].

MAG abundances and assembly statistics were calculated with Mapler (v2.0.1) [58]. Contigs from non-HQ bins and unbinned contigs were concatenated to create 12 pseudogenomes (PG) containing the remaining sequences associated to each phycosphere.

### 2.4 Phylogenomic analysis of *Microcystis*

The phylogenetic placement of the 12 *Microcystis* strains was inferred using a dataset of 349 genomes and method previously described in [34, 57]. Briefly, after predicting the coding sequences of 361 *Microcystis* genomes (including the 12 strains) using Prodigal (v2.6.3), an alignment of 779 concatenated single-copy genes shared by all strains in the analysis (674,919 positions) was generated by Roary (v3.13.0, default parameters except for a protein identity threshold ≥ 90%). Spurious and misaligned regions were removed using trimAl (v1.5) before phylogenetic inference was computed by RaxML (v8.2.12) with the following parameters: GTRGAMMA model, 100 bootstraps. In addition, the average nucleotide identity (ANI) was calculated between each pair of strains in the tree using fastANI (v1.33) [43] to define *genospecies* (ANI ≥97%, [9]), and *genotypes* (ANI ≥99%). Details about the tree construction are available in Supplementary Material and Methods.

### 2.5 Gene prediction and functional annotation

Gene prediction of MAGs and PGs was performed with Prodigal (v2.6.3) [41]. Functional annotations were obtained with eggNOG-mapper (v2.1.12) [39], relying on eggNOG database (v5.0.2) [11], and diamond for sequence alignment (v2.1.9) [8].

Biosynthetic Gene Clusters (BGC) were identified using antiSMASH (v7.0.0) [6] with default parameters. Additional targeted annotations were curated using reciprocal blast searches based on known *Microcystis* metabolic capabilities, as explained in Supplementary Material and Methods. Functions related to biogeochemical cycles were assessed using METABOLIC (v.4.0) [87].

### 2.6 Genome-scale metabolic network reconstruction and comparisons

Genbank files were generated with emapper2gbk (v0.3.1) [4] from gene prediction and functional annotation. Genome-scale metabolic networks (GSMNs) were reconstructed from annotated genomes using Pathway Tools (v27.0) [47] via the mpwt pipeline (v0.8.6) [4]. PADMet library (v5.0.1) [2] was used to manipulate GSMNs and create SBML files [38]. This library was also used to get a description of the GSMNs (*i.e.* size, reaction presence, pathway completeness), and for the curation and integration of missing reactions. For the latter process, we first used the double-blast procedure described in Supplementary Material and Methods for protein annotation, then drafted the reactions catalysed by the identified proteins for integration in corresponding GSMNs with PADMet.

Regarding the pseudogenomes, a single GSMN was reconstructed from concatenated unbinned contigs and low-quality bins per phycosphere, retaining only gene-associated reactions.

Details of GSMNs comparison are described in Supplementary Material and Methods. GSMNs comparisons and metabolome-to-network mapping were performed using MetaNetMap (v1.0.1) [62] (details in Supp. Material and Methods).

### 2.7 Metabolic Modelling

Community metabolic potential was assessed using the Metage2Metabo (v1.6.1) [4] pipeline integrating MisCoTo (v3.2.0) [33] and MeneTools (v3.4.0) [2], based on network expansion (a Boolean abstraction of metabolic producibility) from seed metabolites corresponding to BG11 medium composition (Table S1) to assess reachability of metabolites [29] (details in Supp. Material and Methods).

Two modelling scenarios were explored: i) intra-phycosphere metabolic complementary under laboratory culture conditions, and ii) inter-phycosphere metabolic complementary simulating pond-like environmental conditions (details in Supp. Material and Methods).

### 2.8 Metabolomic data acquisition

Intracellular metabolites from the 12 phycosphere cultures were collected by centrifugation (3,200 g 15°C, 10 min), lyophilised and extracted using complementary monophasic (MP) [28], mainly dedicated to capture specialised metabolites, and biphasic protocols (BP) [59], to maximize metabolome coverage. Details about the MP and BP protocols are provided in Supplementary Material and Methods.

### 2.9 Metabolomic annotation and chemometrics

Feature extraction and annotation of metabolomic signals were performed according to two different workflows. At first, for MP extracts, the mass spectrometry (MS) data were processed using Meta-boScape (v4.0) software (Bruker, Bremen, Germany) in order to generate a data matrix containing semi-quantification results for each analyte in all analysed samples (see details in Supp. Material and Methods). Then, MS-DIAL (v5.25) [79] and MS-cleanR were used to process and clean both MP and BP data and interrogate FragHub spectral libraries [21]. SIRIUS software (v6.1) [27] was used to interrogate structural libraries (SIRIUS database and cyanoMetDB [45]). Output files from MS-DIAL and SIRIUS were further imported into MS-Net [32] in order to suppress analytical redundancy, provide annotation confidence level and classification (NPclassifier, [48]). Annotation workflow parameters are provided in Supplementary Material and Methods. Following MS-Net, spectral library matches were stratified as level 1 (spectral cosine *>* 0.95 and Δ*RT <* 0.2 min), level 2a (cosine *>* 0.85) or 2b (0.7 ≤ cosine ≥ 0.85). *In silico* candidates were supplemented with taxonomic information by querying COCONUT 2.0 [15] using InChiKey-based matching. Among the candidates per feature, those matching taxonomic criteria (identified in bacterial genus or in *Microcystis*) were classified at level 3a and the remaining *in silico* matches stayed at level 3b. Spectral library matches with cosine *<* 0.7 or significant precursor mass errors were classified as level 4 (MS/MS analogues).

Prior chemometrics, replicates were combined through averaging as to keep one signal per metabolite for each phycosphere. Feature intensity was normalised, transformed and scaled (*i.e.* sum normalisation, cube root transformation and pareto scaling). A principal component analysis (PCA) and a hierarchical clustering analysis (HCA) were performed and annotated metabolites were summarised according to chemical classes (NPC classification [48]).

### 2.10 Statistical analysis and data visualisation

Data analysis were handled with python (v.3.12.3), pandas (v.2.2.2), numpy (v.2.1.0), or scikit-learn (v.1.6.1), as well as R (v.4.4.1) package vegan (v.2.6 6.1). Visualisation was generated using python matplotlib package (v.3.9.2), and R package dendetextend (v.1.17.1) for tanglegrams.

Phycosphere taxonomic trees were constructed in newick format using GTDB-tk results [67], and displayed with the iTOL tree display and annotation tool [52]. GSMNs similarity across phycospheres were assessed by performing KR Clarke analysis of similarity [19], and visualised with Principal Coordinates Analysis (PCoA). For the latter, we considered the PCoA satisfactory if the condition *stress* − 1 *<* 0.1 was respected, according to [50].

## 3 Results

### 3.1 Phylogenomic and potential metabolism of the 12 *Microcystis* strains

A total of 9,006,345 HiFi metagenomic reads (67.9 Gbp) was obtained, ranging from 516,628 and 1,021,017 per phycosphere community, with median contig lengths between 5,973 to 7,951 bp. One complete *Microcystis* genome (single contig) was reconstructed from each non-axenic culture, with an average size of 5.3 Mbp.

#### 3.1.1 Phylogenomics of *Microcystis* strains

*Microcystis* phylogeny was inferred from a concatenated alignment of 779 single-copy genes (674,919 aa positions) shared by 361 strains. The twelve strains captured local *Microcystis* diversity and were distributed across the phylogenomic tree, with *Microcystis* sp. PMC1371.22 and *Microcystis* sp. PMC1372.22 closest to the root (Fig. 1A). Using ANI threshold ≥ 97% [9], 21 genospecies (A-U) gathering multiple strains were identified encompassing 352 strains. The twelve strains belonged to six distinct genospecies (C, D, F, I, S, U). All strains within a genospecies belong to a single genotype (ANI ≥ 99%)), except for genospecies I, which three strains belong to three distinct genotypes.

**Figure 1:**
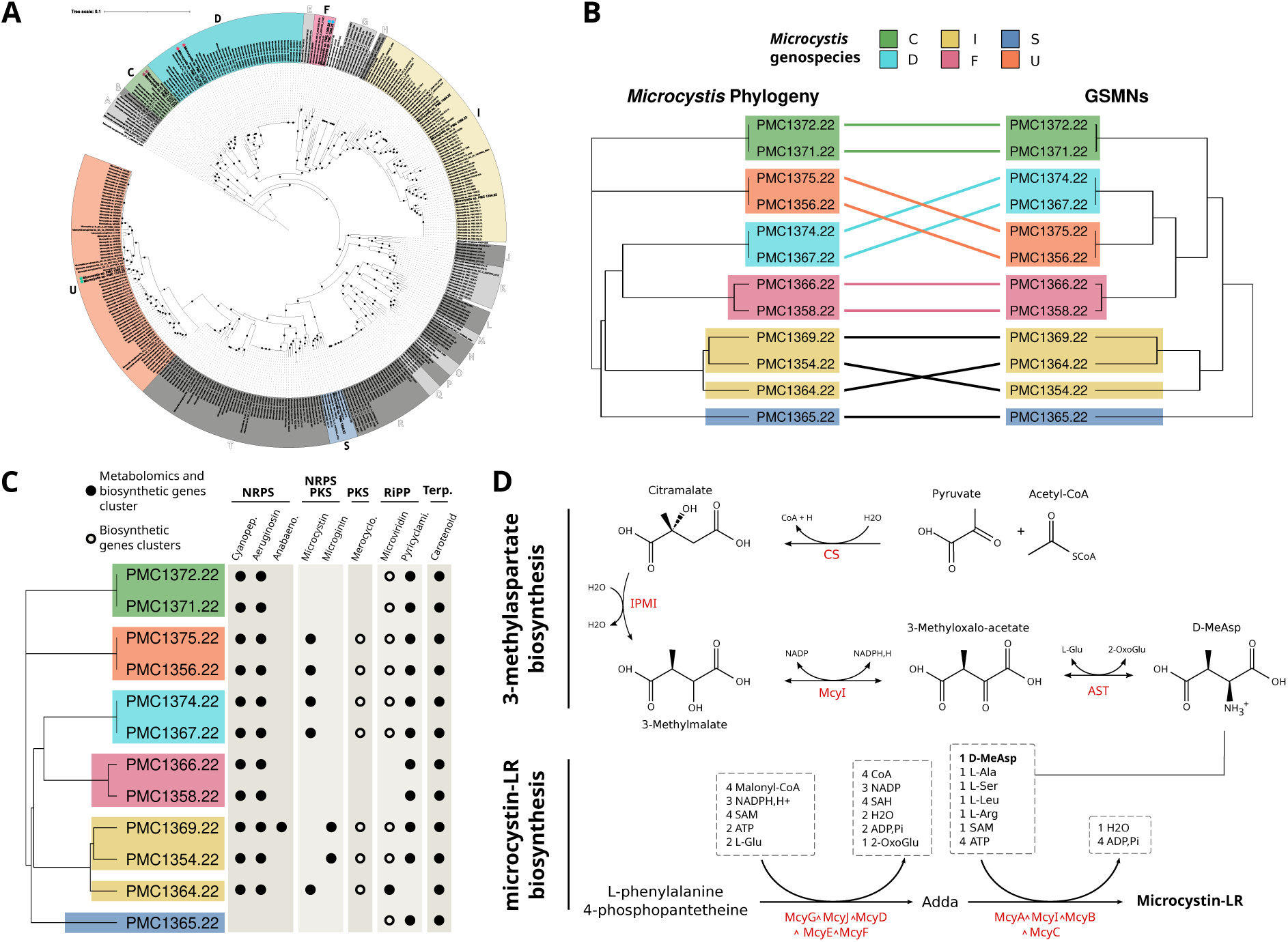
Reconstruction of metabolism for the 12 *Microcystis* strains. **A.** Maximum likelihood phylogenomic tree of *Microcystis* species. The boxes group together strains from the same genospecies, labelled from A to U. The coloured ones (C, D, F, I, S and U) correspond to the genospecies of the surveyed strains (in bold). The stars associated with the labels indicate that the associated strains belong to the same genotype. Tree branches supported by bootstrap values over 60% are indicated by a black circle. The tree is rooted on *M. aeruginosa* Ma SC T 19800800 S464, as determined in previous studies [9]. **B.** Tanglegram of *Microcystis* ANI phylogenetic tree on the left, and of the hierarchical clustering of GSMN’s reactions presence matrix on the right. Hierarchical clustering was conducted using Jaccard distance and complete-linkage clustering. Species on branches coherent between the two dendrogram are connected by a coloured link. **C.** Specialised metabolism of *Microcystis* strains. On the left, the phylogenetic tree of *the 12 Microcystis* strains. On the right, matrix of the specialised metabolites biosynthetic genes cluster (BGC) presence in genomes, and metabolite identification in the phycosphere according to metabolomics from multiplexed LC-HRMS/MS analyses (Supp. Material). **D.** Microcystin-LR and 3-methylaspartate biosynthesis, according to literature and identified genes in the twelve *Microcystis* genomes. At the bottom, reactions of the MC-LR biosynthesis are presented as integrated in the GSMNs, that is as two lumped reactions. **Abbreviations: Anabeno**., anabenopeptin; **Cyanopep**., cyanopeptolin; **Merocyclo**., merocyclophane; **NRPS**, non-ribosomal peptides synthase; **PKS**, polyketide synthase; **Pyricyclami**., pyricylamide; **RiPP**, ribosomally synthesized and post-translationally modified peptides; **Terp**., terpenoids; 2-OxoGlu, 2-oxoglutarate; **Adda**, (25,35,85,9S)-3-amino-9-methoxy-2,6,8-trimethyl-10-phenyl-4,6-decadienoic, **AST**: aspartate transaminase; **CS**, citramalate synthase; **D-MeAsp**, D-methylaspartate; **IPMI**, isopropylmalate isomerase acid.

#### 3.1.2 Potential functional metabolism of *Microcystis*

GSMNs reconstructed for the twelve *Microcystis* genomes were highly similar (from 1,780 to 1,823 reactions; Table 1), consistent with their phylogenetic proximity (Fig. 1A). The quality of GSMN reconstruction was validated by the congruence between metabolic distances across strains and GSMN contents (Fig. 1B). This suggests that the reconstruction captured clade-specific metabolic features. *Microcystis* GSMNs were identical within genospecies D (PMC1374.22 and PMC1367.22) and U (PMC1375.22 and PMC1356.22), although undetected differences may exist due to annotation limits, and the prevalence of unknown cyanobacterial genes [12].

**Table 1:**
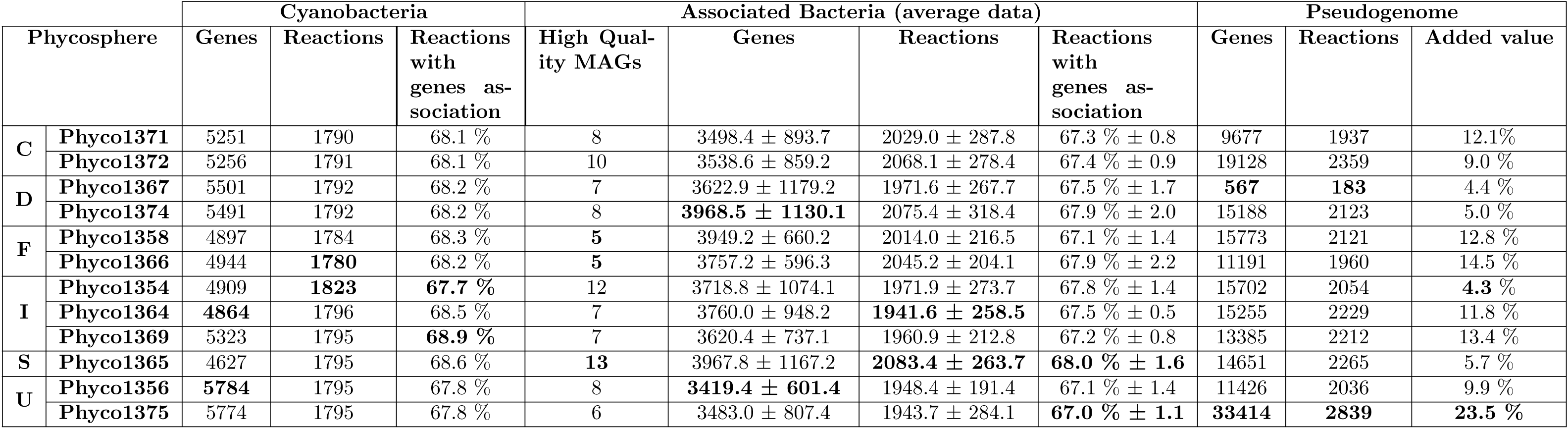
Description of the twelve phycosphere metagenomes and their reconstructed GSMNs. For each community, a distinction is made between *Microcystis*, its associated bacteria and the pseudogenome (PG) that is reconstructed from the remaining contigs of the assembly. PG’s reactions introduced in the phycosphere is the percentage of reactions of the pseudogenome’s metabolic network that are not in any of the other phycosphere’s members GSMN. On the very left column and coloured, the *Microcystis* genospecies is indicated. Bold: minimum and maximum values.

Specialised metabolism (SM) largely encoded by biosynthetic gene clusters (BGCs) was surveyed across genomes (Fig. 1C, Supp. File 1). BGC content was strictly conserved within genotypes and more loosely conserved within genospecies [40], as illustrated by genospecies I, which exhibits greater genetic divergence (97% *<* ANI *<* 99%) among its three strains belonging to distinct genotypes. Only five strains from genospecies U, D and I carried microcystin-associated BGCs while carotenoid-associated BGCs were detected in all strains. *Microcystis* PMC1365.22 (genospecies S) lacked most BGCs. Most SM products were detected in intracellular metabolomes, except microviridin, detected only in PMC1364.22, potentially due to sequence variability and annotation limits.

SM-associated reactions were absent from GSMNs and the MetaCyc database [13] highlighting the limited representation of SM in GSMN reconstruction as previously established [73]. To address this, the microcystin-LR biosynthetic pathway was manually curated (Fig. 1D), integrating literature (Sup. Table S8) and BGCs annotations. Overall, *Microcystis* strains shared a conserved core metabolism with phylogenetically structured SM diversity, including variation in pigment and toxin production.

### 3.2 Taxonomy and functions of the associated microbiome

#### 3.2.1 Taxonomic composition of phycosphere’s associated bacteria

Ninety-six high-quality (HQ) MAGs (5-13 per phycosphere) of associated bacteria (AB) were recovered, including 62 MAGs nearly, or fully complete and assembled in a unique contig (Supp. File 1). HQ MAGs accounted for *>* 90% of read abundance in most phycospheres, with *Microcystis* genomes showing the highest coverage (82X to 1,115X) consistent with culture conditions (Supp. Fig. S1). Cyanobacterial relative abundance varied among phycospheres, with *Phyco1365* (associated to *Microcystis* PMC1365.22), *Phyco1356* (associated to PMC1356.22) and *Phyco1375* (PMC1375.22) displaying lower dominance, suggesting distinct community structures (Fig. 2A, Supp. Fig. S1). Unbinned contigs and lower quality MAGs were merged into phycosphere-specific pseudogenomes (PGs) to capture residual functional diversity from low-abundant unassembled or unbinned populations.

**Figure 2:**
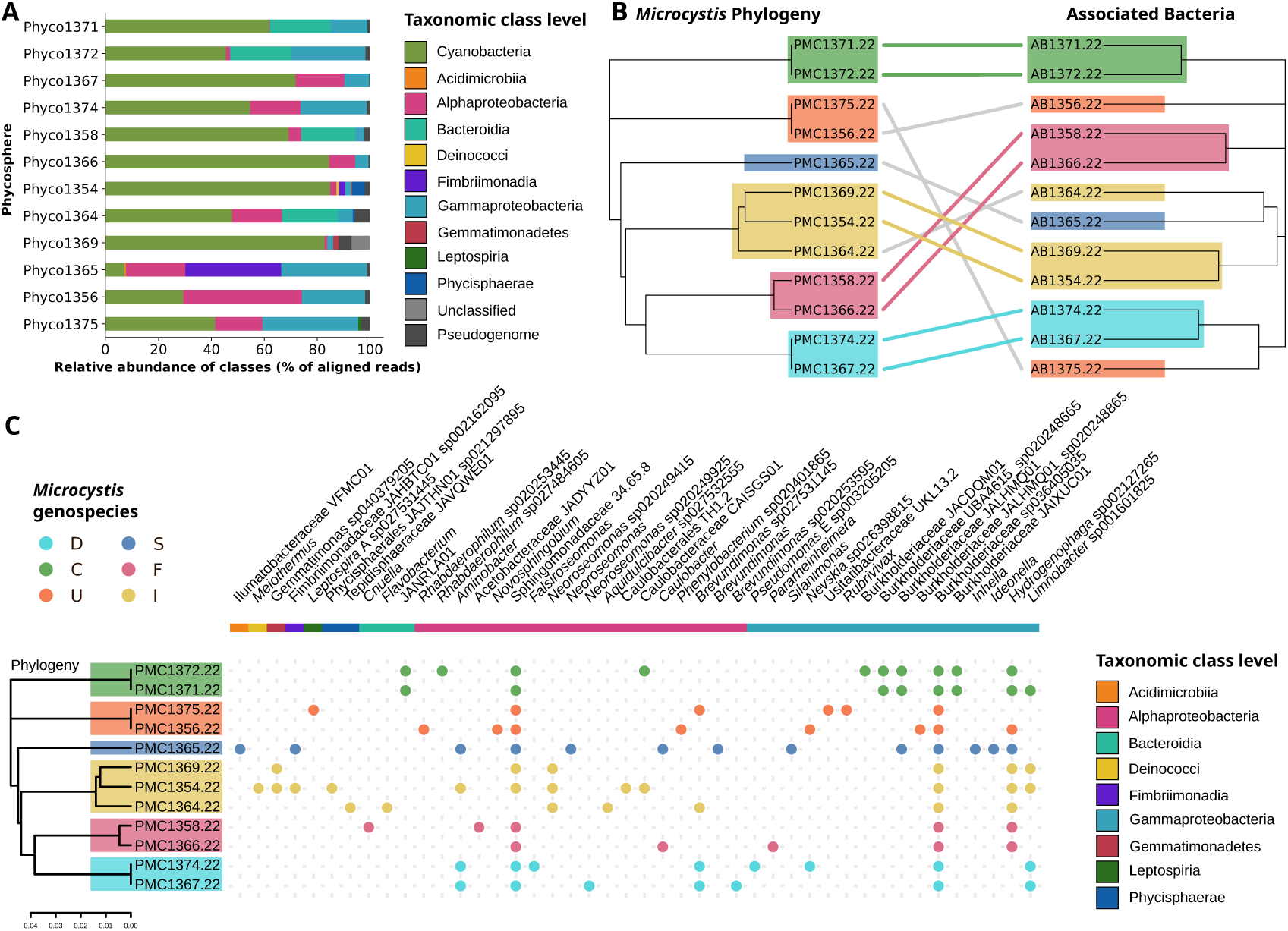
Metagenomic assembly of the 12 phycospheres. **A.** Class taxonomy level and relative abundance of the associated bacteria based on metagenomic read mapping **B.** Relationships between *Microcystis* strain phylogenomy and the taxononic composition of the respective associated bacteria according to hierarchical classification **C.** Heatmap of the bacterial MAGs occurrence within the twelve phycospheres.

Taxonomic assignment of MAGs (Fig. 2A, Supp. Fig. S2) revealed that most AB belonged to the *Alphaproteobacteria*, *Gammaproteobacteria*, or *Bacteroidia*. Species-level composition of phycospheres (Fig. 2C) often mirrored *Microcystis* phylogeny, with several MAGs associated with specific genospecies: *e.g. Neoroseomonas sp020249415* with genospecies I, *JANRLA01* (Bacteroidia) and *Burkholderiaceae UBA4615 sp020248665* with genospecies C. In contrast, *Burkholderiaceae sp036405035* and *Sphingomonadaceae 34.65.8* were nearly ubiquitous, indicating stable associations across genospecies. *Fimbriimonadia* were enriched in *Phyco1365* which showed lower cyanobacterial dominance, and also found in smaller proportion in *Phyco1354*.

Phylosymbiosis [72] was assessed by comparing the phylogeny of *Microcystis* with the dissimilarities of the bacterial community (Fig. 2B). The strongest congruence occurred at species and genus levels (Normalised Robinson-Foulds (NRF) metric [75]: 0.55), with clustering by genospecies for several genotypes (D, F, C, and I (2/3 strains)). The dendrogram of genospecies U and its AB was not obviously consistent between the two distinct strains, according to slight AB community variation. Higher taxonomic ranks (family and order) showed weaker strains specificity and congruence (NRF: 0.88 and 0.77 resp. Supp. Fig. S3), indicating that phylosymbiosis may primarily operate at fine taxonomic resolution, that would imply even more pronounced metabolic specificities.

#### 3.2.2 Metabolic potential of phycosphere’s associated bacteria

GSMNs of AB and PGs were heterogeneous and generally larger than *Microcystis* networks, averaging 2,010 reactions (Table 1). Some PGs yielded small networks (*e.g. Phyco1367*) whereas others were larger but largely redundant with the rest of the phycosphere. PGs of *Phyco1366* and *Phyco1375* contained many genes and brought the most additional unique reactions to the community, with respectively 14.5% and 23.5% of unique reactions, suggesting the presence of low-abundance - uncaptured - populations in those phycospheres with distinct metabolic capabilities.

To further investigate functional redundancy and complementarity within phycosphere communities, we performed a rarefaction analysis of the number of enzyme commissions (EC) and other functional annotations in *Microcystis* and AB genomes, excluding PGs (Fig. 3A, Supp. Fig. S4). The absence of a plateau indicates that even in these small cultivated phycospheres, each member likely contributes unique funtions. However, the curve inflected beyond five members, suggesting diminishing functional novelty with increasing community size. The union of all reconstructed phycosphere genomes set an upper limit of ∼ 300 additional EC numbers, whereas PGs contributed ∼ 100 EC numbers absent from other genomes. Analysis of genes related to biogeochemical cycles (Supplementary Material) revealed clear functional partitioning between community members. Cyanobacteria and AB differed in nitrogen metabolism, whereas heterotrophic bacteria encoded pathways for the degradation of complex carbon substrates.

**Figure 3:**
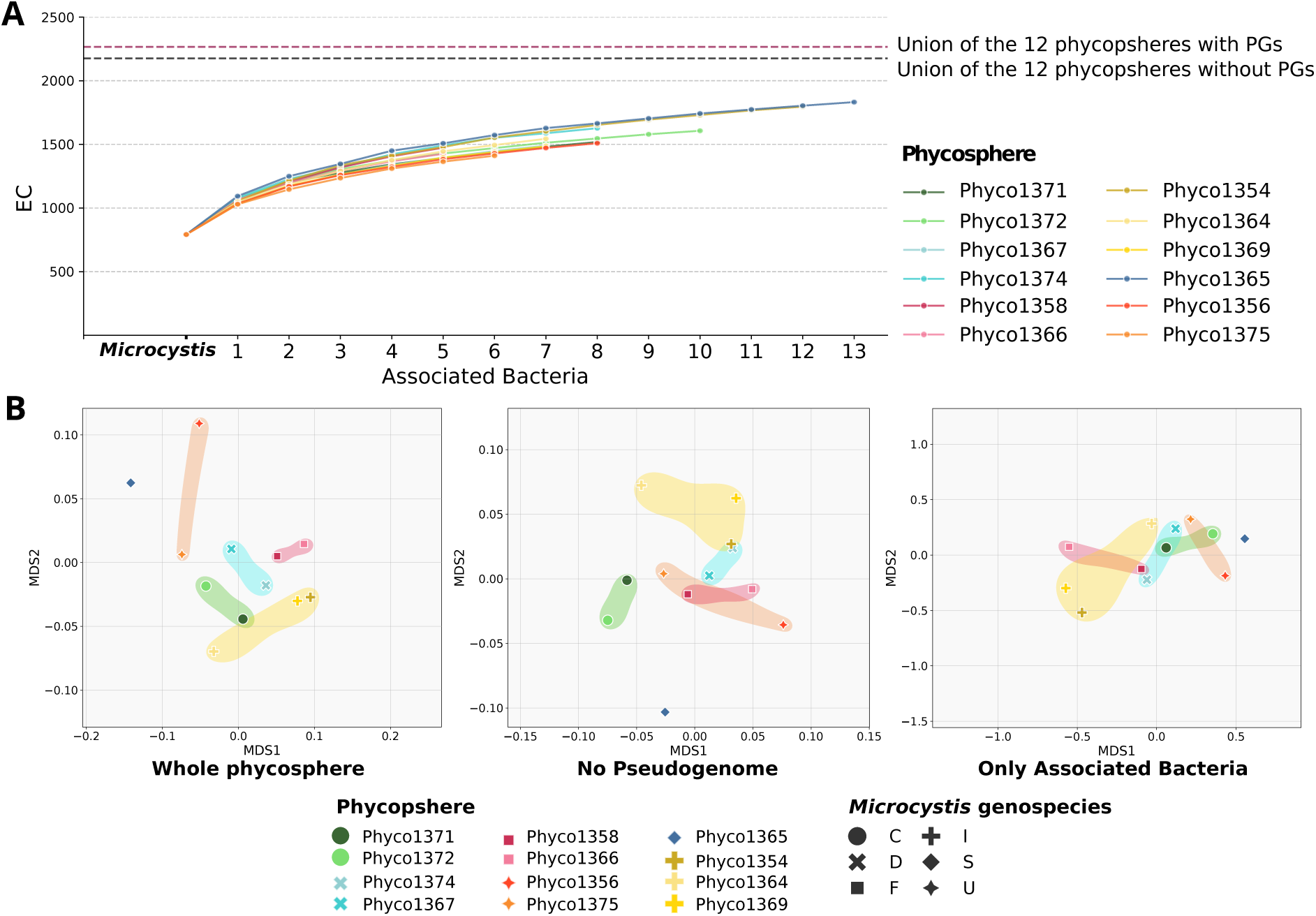
Metabolic complementarity within phycospheres. **A.** Rarefaction curve of EC numbers annotated in phycosphere metagenomes. Rarefaction curves represent the accumulation of non-redundant EC numbers as a function of the number of sampled associated bacteria, based on 100 sampling iterations. The starting point is *Microcystis* EC numbers. PGs are excluded from the sampling of phycosphere’s individual curves. **B.** Principal Coordinates analyses of the GSMNs reactions content, weighted by species abundance and grouped by phycosphere. From left to right; for the whole phycospheres, without the pseudogenome, and with only the associated bacteria. Shaded areas show phycospheres associated to *Microcystis* strains belonging to the same genospecies. **Abbreviations:** EC, enzyme commission number; PG, pseudogenome

We next compared metabolic reaction repertoires across phycospheres using three configurations: all members including PGs, all members excluding PGs, and AB only (Fig. 3B). Complete phycospheres showed a significant but moderate clustering according to *Microcystis* genospecies (KR Clarke analysis of similarity [19], *R* = 0.539, p-value= 0.007). This pattern persisted when PGs were excluded (*R* = 0.554, p-value= 0.005) but weakened when only AB were considered (*R* = 0.317, p-value= 0.084). In all analyses, phycospheres associated with genospecies I displayed the highest functional diversity.

### 3.3 Meta-metabolomes of the phycospheres

The merged endometabolome features (from both MP and BP extractions; Supp. Fig. S5, Fig. 4A) of the 12 phycospheres comprised 20,931 features: 12,608 with only MS1 and 8,323 with MS2 spectra. Among these, 0.6% (135 features) and 0.3% (63 features) were annotated into levels 2a and 2b, respectively. Including *in silico* candidates (level 3a, 144 features and 3b, 5674 features), up to around 31% of features were putatively annotated. Both HCA (Fig. 4A) and PCA (Supp Fig. S6) revealed clustering of phycospheres according to *Microcystis* genospecies, consistent with a massive cyanobacterial-driven metabolite production and likely influenced by high *Microcystis* dominating biomass within cultures, that is also fully congruent with our metagenomic observations considering the above 1000-times higher biovolume of *Microcystis* cells compared to heterotroph AB (Fig. 2A). Only slight deviations were observed with *Phyco1369* diverging from genospecies I and *Phyco1365* showing a metabolomic profile closer to genospecies U (Fig. 4A). Annotated metabolites (6,016 features with L2 or L3 confidence) were categorised by NPC classification (Supp. Fig. S7) and the 20 most represented superclasses were highly similar across phycospheres (Fig. 4B). Only 144 of the annotated features could be specifically linked to taxa detected by metagenomics, mainly *Microcystis*, *Pseudomonas*, and *Burkholderiaceae* (details, including putative ecological roles and compound identities, in Supp. Material, Supp. Fig. S8).

**Figure 4:**
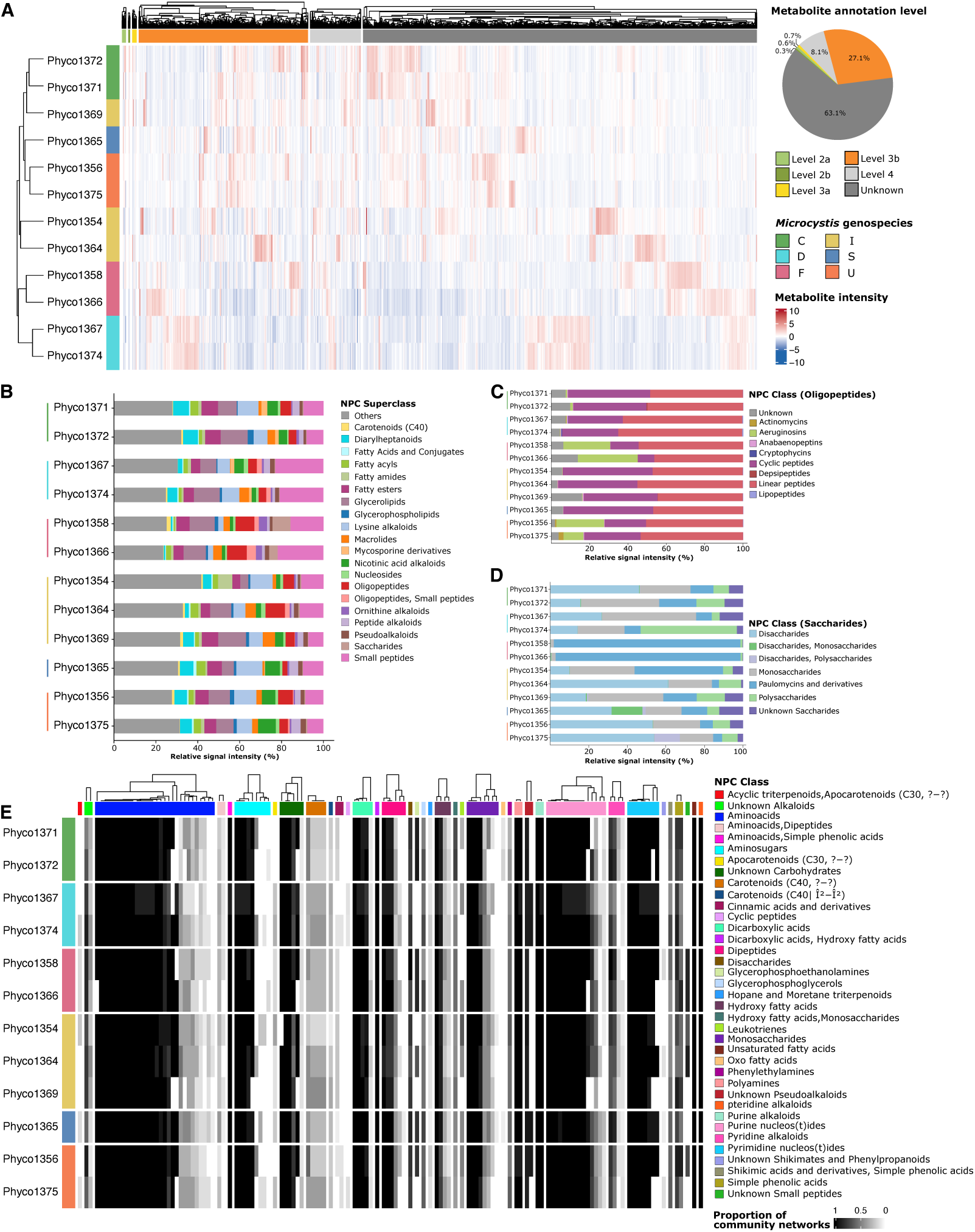
Metabolomic analysis of the 12 phycospheres. **A.** Hierarchical clustering analysis on phycospheres transformed endometabolome signals. For each phycosphere, triplicates were averaged to merge the different extractions results and get one set of transformed signals per community. Metabolites annotations was conducted as previously described, and evaluated with levels assigned as described in Methods. Hierarchical clustering was based on euclidean distance and complete linkage method. **B.** Barplot depicting the relative abundance of L2 and L3 annotated metabolomic signals in phycospheres stratified by NP Classifier superclasses. Only the 20 most abundant superclasses are depicted, others being aggregated in *Others*. **C.** Focus on the annotated metabolomic signal classified as “Oligopeptide” superclass (relative abundance). The 20 most abundant classes are depicted, others being aggregated in *Others*. Cyclic peptides include microcystins, additional toxins and other specialised metabolites, such as aeruginosins, or cyanopeptolins. **D.** Focus on the annotated metabolomic signal classified as “Saccharides” superclass. Relative abundance of Saccharides classes in the twelve phycospheres. **E.** Heatmap depicting the mapping of L2-L3 annotated metabolomic signals onto the GSMNs of all phycospheres members. Heatmap values represent the proportion of GSMNs per phycosphere that contain the related metabolite. Identification in GSMN means the metabolite is a substrate or a product of a metabolic reaction. Metabolites are stratified by their NP-Classifier classes. Cyclic peptides correspond to Microcystin-LR. Carotenoids include echinenone, canthaxanthin, zeaxanthin, adonixanthin and dike-tospirilloxanthin. **Abbreviations:** NPC, NP Classifier.

The oligopeptide NPC superclass (14% of annotated compounds), showed strong variability among phycospheres, with genospecies-level similarities (Fig. 4C). Actinomycins and linear-lipopeptides were specific to genospecies U and C, respectively, while microcin SF608, likely produced by AB, was enriched in genospecies U. Microcystins (e.g. microcystin-RR, MC-LR) were most abundant in *Phyco1374* and *Phyco1367* (genospecies D), *Phyco1375* and *Phyco1356* (genospecies U), and especially in *Phyco1364*, matching BGC-based predictions (Fig. 1C). Additionally, a few low-intensity putative microcystins were detected in strains lacking microcystin BGCs. These are likely false positives arising from limitations of automated metabolite annotation, as certain micropeptins/cyanopeptilins share structural features with microcystins but can be manually distinguished based on the presence of specific diagnostic ions.

Saccharides, for their potential role in cross-feeding [61], were examined and varied by genospecies (Fig. 4D); for example, paulomycins were most abundant in genospecies F phycospheres, and some monosaccharides were detected exclusively in *Phyco1365*, the sole representative of genospecies S (Fig. 1C, 2A). No relevant metabolites were detected in the exometabolome beyond culture medium components, suggesting a rapid consumption of extracellular components by heterotroph AB.

Mapping metabolomic features to phycosphere GSMNs yielded only 133 matches (Fig. 4E). Mapped metabolites were mainly amino acids, amino sugars, dipeptides, purines, pyrimidines, and nucleosides, with variable distribution across GSMNs. Notably, carotenoids were mapped only to GSMNs of AB, including PGs, that may correspond to a false negative regarding *Microcystis* which carotenoid biosynthetic genes remain poorly characterised and further automatically annotated.

### 3.4 Modelling the intra- and inter-phycosphere metabolic complementarity

To investigate metabolic complementarity within and across phycospheres, we simulated metabolite production for each GSMN under BG-11 medium conditions, *i.e.* we identified the metabolites of the GSMNs that could be reached from these nutrients. Predictions distinguished metabolites produced individually from those requiring mutualistic interactions. We tested two scenarios: (i) separate simulations for each phycosphere to assess within-community complementarity, and (ii) a merged network simulation to predict larger-scale interactions between phycospheres (Fig. 5)

**Figure 5:**
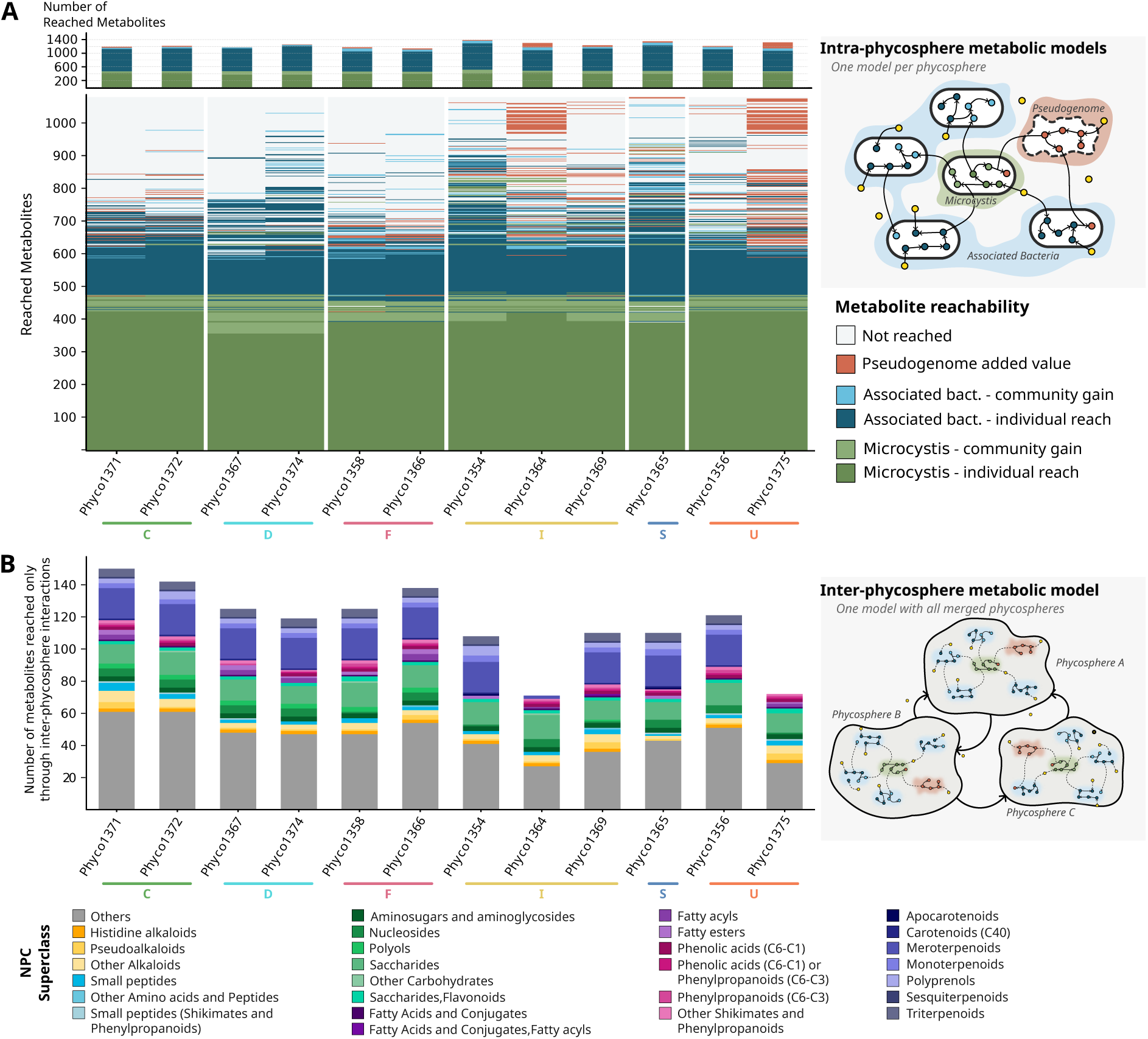
Modelling the metabolic complementarity intra- and inter-phycospheres. **A.** Heatmap of metabolites reach across each phycosphere metabolic capabilities modelling, grouped by genospecies (C, D, F, I, S, and U). Producers of reachable metabolites were grouped to distinguish the roles of *Microcystis*, associated bacteria (AB), and pseudogenomes (PGs), as well as to assess the impact of interactions within and between these groups. Each row is a metabolite, preferably shown as reached by the cyanobacteria, when it is also reached by one of the AB. Non-overlapping reachability is presented in Supplementary Figure 9. “Individual reach”, metabolites produced by at least one community member in the phycosphere; “Added value”, metabolites reached only through metabolites exchanges; “not reached”, metabolites not reached in modelling or absent from GSMNs. The upper barplot enumerates the (possibly overlapping) metabolites reached in each of the observed situation (by *Microcystis*, AB, through the pseudogenome). **B.** Classification of inter-phycosphere added-value. For each phycosphere, metabolites that are reached by one of their member only through phycosphere-phycosphere exchanges are classified according to their NPC Superclass level.

Intra-phycosphere metabolic potentials and interactions are shown in Fig. 5A. Under the simulated conditions, *Microcystis* could access a core of over 400 metabolites, largely autonomously. Additional metabolites became accessible only through metabolic interactions with AB and PGs, with variability among phycospheres. Differences among cyanobacterial GSMNs were largely offset by community composition (Fig. 1, Table 1). In contrast, AB metabolic potential varied despite a core of 510 shared metabolites and did not align cyanobacterial genospecies (analysis of similarity [19], R=0.355, p=0.07). PGs unlocked additional pathways, contributing approximately 25 unique metabolites per phycosphere. *Phyco1364* and *Phyco1375* shared unique predicted cyanobacterial metabolites, mainly terpenoids. Comparisons between model predictions and metabolomics are detailed in Supp. Material.

Inter-phycosphere simulations predicted access to approximately 100 additional metabolites per community (Fig. 5B), including fatty acids, shikimates, phenylpropanoids, carbohydrates, and terpenoids. Notably, terpenoid functions from *Phyco1364* and *Phyco1375* spread to other communities, suggesting transferable metabolic benefits.

Together, these results demonstrate that metabolic complementarity enhances metabolite accessibility both within and between phycospheres and is shaped by phycosphere composition.

## 4 Discussion

### *Microcystis* metabolism reflects genospecies structure

*Microcystis* genomes are highly plastic [12], with non-ribosomal peptide synthase and polyketide synthase gene clusters accounting for 5% of the genome [78], and up to 240 natural products classified as specialised metabolites through genomics [51, 25]. Given their high versatility, capturing metabolic variation across genospecies is therefore essential. Our GSMN reconstructions revealed a robust consistency between metabolic reaction content and genospecies phylogeny. BGCs repertoires were similarly conserved within genospecies, except for genospecies I which displayed greater heterogeneity. Metabolomic analyses further confirmed the production of multiple microcystin variants by *Microcystis*. However, their biosynthetic pathways were largely absent from GSMNs due to incomplete database representation. Manual curation of the MC-LR pathway in GSMN underscores the substantial efforts still required to incorporate SM into genome-scale models and could benefit from existing efforts in BGC description [73, 85].

### Phycospheres harbour a reduced taxonomic diversity driven by *Microcystis* phylogeny

The reduced taxonomic diversity observed in phycospheres is consistent with culture conditions [88, 49, 37], and reflects selective pressure favouring *Microcystis* growth thus prioritising AB that are essential for it, or at least non-antagonistic [49, 83]. It was previously reported that bacterial communities in the phycosphere are shaped by the genotypic characteristics of *Microcystis* [72]. Despite the limited dataset, this observation implies that bacterial partners conserved through cultivation process may present specific selective added-value that directly depends on *Microcystis* genospecies. The molecular mechanisms behind, potentially mediated by metabolic complementary or selective processes, remain however unknown so far.

Dominant taxa, including Caulobacterales and Burkholderiales, are frequently reported in cyanobacterial phycospheres in both environmental and lab studies. The ubiquitous presence of *Hydrogenophaga* sp. aligns with its recurrent association with *Microcystis*, in natural environments and lab cultures [49, 54, 88], while *Sphingomonadales*, known for microcystin tolerance and degradation [24, 81], were systematically detected. While the metabolomic and BGC analyses confirmed microcystin production within the phycospheres, the absence of known microcystin degradation genes (*mlr*) suggests that tolerance rather than active degradation, may be the main adaptation mechanism of these bacterial partners.

### Functional complementarity within the phycosphere, but a cyanobacteria-driven metabolome

Comparative functional analyses and GSMNs revealed substantial diversity and metabolic complementarity among phycosphere members. Notably, PGs contributed up to 23% unique metabolic reactions to phycosphere, highlighting the underestimated role of low-abundance community members. While AB functions alone did not reflect *Microcystis* genospecies, genospecies-specific signals emerged when considering the full phycosphere. *Microcystis* strains likely influence their communities via their own metabolism [72] but the lack of a clear signal at the global AB genomic level suggests that community shaping depends more on specific metabolic functions or environmental responses. Functional complementarity included nitrate and nitrite reduction by AB, potentially supporting redox balance and nutrient recycling [54, 37]. In contrast, thiosulfate oxidation genes were absent from *Microcystis* but present in *Limnobacter* spp. in half of the phycospheres, consistent with prior detection during bloom events [54]. Yet, metabolism captured by metagenomics and annotation likely reflects only a fraction of the total metabolic landscape, as illustrated by the limited fraction of annotated metabolites that could be directly linked to GSMNs, highlighting opportunities for improvement [31].

Metabolomic annotations captured only a small fraction of all detected signals, with fewer than 2% annotated at level 2 and most assigned *in silico*. The absence of annotated exometabolites prevented formal inference of metabolite exchange, possibly due to optimal growth conditions or rapid metabolite turnover. Nonetheless, endometabolome analyses demonstrate the value of combining multiple extraction and acquisition strategies to reveal chemical diversity beyond what metagenomics alone can capture.

Endometabolomic profiles mirrored *Microcystis* phylogeny, reflecting cyanobacterial dominance in biomass and metabolites production. Most annotated compounds belonged to small peptides, glycerolipids, and oligopeptides, which discriminated phycospheres. The tight coupling between *Microcystis* genospecies and chemical profiles aligns with recent reports linking genotypes and chemotypes in environmental phycospheres [40]. Among the phycosphere-specific chemical signatures, we surveyed the oligopeptide and saccharide signals. Cyanobacterial oligopeptides, known for their structural diversity and inhibitory activities, likely contribute to metabolic plasticity and defence [1, 63, 7]. Saccharide analyses revealed the importance of paulomycins, particularly in genospecies F, although their typical producer *Streptomyces* [53] was absent from high-quality MAGs. This may reflect low-abundance or uncharacterised producers, or limitations of *in silico* metabolite annotation.

*Phyco1365* deviated from others with distinct BGC content, taxonomic structure, phycosphere GSMNs, and metabolomic profiles. Differences in endometabolome content are likely due to low cyanobacterial abundance, distinct AB, and the near-exclusive presence of *Fimbriimonadia*, the latter being also observed at a lower abundance in *Phyco1364*, the second endometabolome outlier. These differences suggest altered microbial interactions but may also reflect the incompleteness of reference databases.

### Ecological insights from cultivated freshwater phycospheres

Recent studies have shown that metabolomic profiles of environmental *Microcystis* strains closely reflect their genotype, likely shaped by local environmental conditions [40]. Although our study relies on cultivated phycospheres, our results support this genotype–chemotype link and its ecological relevance. While Huré et al. [40] identified 13 chemotypes from 65 strains collected from 13 lakes, here we detected six distinct genotypes within a single site, indicating that multiple genotypes can coexist under similar environmental conditions and that AB patterns may persist under lab-cultivation.

Several annotated compounds have reported allelopathic or bactericidal properties, including sorbistin A2, ornibactin C6/C8, and paulomycins [80, 60, 53], and may contribute to shaping phycosphere bacterial communities. However, as these assignments rely solely on *in silico* annotation, they should be interpreted with caution.

Regarding toxin production, the inability of some *Microcystis* strains to produce microcystins did not result in major differences in the overall phycosphere metabolism. How microcystin-producing and non-mycrocystin-producing strains interact at the bloom scale remains unresolved, but our modelling approach highlights functional complementarity, supporting potential *phycosphere*–*phycosphere* interactions at fine spatial scales [35]. These interpretations remain limited by incomplete reference databases, meaning GSMN reconstructions provide only a partial view of metabolism [73]. Given the number of analysed genomes, we therefore relied on discrete modelling approaches [29, 4, 3]. Although cultivated phycospheres do not fully recapitulate bloom conditions, they nonetheless reveal substantial diversity and potential interaction networks, offering valuable insights into freshwater microbial ecology.

### Intra- and inter-phycosphere metabolic interactions: ecological perspectives from metabolic modelling

This study explored micro-diversity at multiple organisational levels. We first characterised intra-population genomic micro-diversity within the genus *Microcystis* across twelve naturally co-occurring isolates, consistent with patterns previously reported during bloom events [36, 77]. In parallel, micro-diversity was also observed within the AB communities of each phycosphere, with strong congruence between cyanobacterial genotypes and their bacterial partners. Although isolation and laboratory cultivation reduced community complexity, core *Microcystis*–AB interactions appeared to be preserved across genospecies. Together, these results extend the well-described micro-diversity of *Microcystis* to the phycosphere level, where it is thought to support adaptation to fine-scale ecological niches [36].

Metabolic complementarity simulations further indicated that phycospheres can collectively expand their metabolic repertoire beyond individual organismal capacities. Rather than solely reflecting functional redundancy, this complementarity enables access to otherwise unreachable metabolites and aligns with previous observations of cooperative metabolism in cyanobacteria-associated communities [72]. Maintaining phycosphere micro-diversity at small spatial scales may enhance resource use efficiency by AB and support *Microcystis* growth, contributing to bloom persistence and providing a more relevant framework for modelling bloom dynamics [71].

When modelling interactions among the twelve phycospheres, an additional increase in metabolic potential emerged, highlighting that phycosphere metabolism is shaped not only by *Microcystis*–AB interactions but also by complementary processes operating across neighbouring communities. During bloom events, closely co-occurring phycospheres may thus be viewed as neighbour ecotypes represented by distinct genotypes. We therefore hypothesise that environmental adaptation during blooms should be studied at the metacommunity level, and that maintaining phycosphere micro-diversity through interactions among closely related phycospheres represents a potential ecological advantage.

Overall, the systems biology framework adopted here demonstrates the value of integrating metagenomics, metabolomics, and metabolic modelling to investigate the structure and function of *Microcystis* phycospheres. We show that *Microcystis* phylogeny influences both bacterial recruitment and the chemical landscape, while simulations highlight the potential for metabolic cross-feeding within and across phycospheres. These predictions generate testable hypotheses on mutualistic interactions and resource-sharing strategies in *in situ* metacommunities across contrasting ecotypes that now require experimental validation.

## Supporting information

Supplementary content

## Supplementary information

Supplementary files include the following:

- Supplementary Material *supplementary material.pdf* : additional text, figures and tables related to methods and results
- Supplementary File 1 *Supplementary tables.xlsx* : tables summarising the characteristics of MAGs reconstructed from metagenomic data, and antiSMASH results.

## Data availability

Raw reads and assembled genomes are available on EBI ENA under accession number PRJEB102163. Metabolomic data is available on MetaboLights [84] with the study identifier MTBLS13909 (reviewer access: https://www.ebi.ac.uk/metabolights/reviewerdd864ca1-e28b-44b3-9706-532e6ff40fdc). Processed data related to the analyses are shared on Recherche DataGouv https://doi.org/10.57745/ 2P0IKJ. Scripts to perform data analysis are available on https://gitlab.inria.fr/comic-project/ COMIC-Metabo.

## Acknowledgements

Experiments presented in this paper were carried out using the PlaFRIM experimental testbed (https://www.plafrim.fr), supported by Inria, CNRS (LaBRI and IMB), Université de Bordeaux, Bordeaux INP and Conseil Régional de la Nouvelle Aquitaine. We acknowledge the GenOuest bioinformatics core facility (https://www.genouest.org) for providing the computing infrastructure. We acknowledge the METABOHUB research infrastructure (ANR, 11-INBS-0010, https://www.metabohub.fr/)

## Authors contributions statement

**Juliette Audemard**: methodology, software, formal analysis, investigation, writing - original draft, visualization, writing - review & editing. **Nicolas Creusot**: conceptualization, methodology, supervision, visualization, writing - review & editing, project administration, funding acquisition. **Julie Leloup**: resources, validation, writing - review & editing, supervision, funding acquisition. **Charlotte Duval**: resources. **Sébastien Halary**: resources, data curation, review & editing. **Lou Mary**: formal analysis, visualization, review & editing. **Mélissa Eon**: resources, review & editing. **Thomas Forjonel**: resources. **Mohamed Mouffok**: resources, formal analysis, investigation. **Rémy Puppo:** formal analysis **Elodie Belmonte**: resources, writing, review & editing. **Véronique Gautier**: resources, writing, review & editing. **Jeanne Got**: writing - review & editing. **Marie Lefebvre**: writing - review & editing. **Gabriel V. Markov**: formal analysis, writing - review & editing. **Coralie Muller**: formal analysis, software. **Benjamin Marie**: validation, writing - original draft, writing - review & editing, supervision, funding acquisition. **Binta Diémé**: conceptualization, formal analysis, investigation, methodology, supervision, writing - review & editing, project administration, funding acquisition. **Clémence Frioux**: conceptualization, formal analysis, investigation, methodology, writing - original draft, visualization, supervision, writing - review & editing, funding acquisition.

