## Supplementary content for "Integrating metagenome-scale metabolic modelling and metabolomics to identify biochemical interactions in *Microcystis* phycospheres"

#### 1 Supplementary material and methods

##### 1.1 Metagenomic sequencing

Single-molecule Real-time long reads sequencing was performed at Gentyane Sequencing Platform (Clermont-Ferrand, France) with a PacBio Revio Sequencer (Pacific Biosciences, Menlo Park, CA, USA). A library was prepared using the SMRTbell® Template Prep kit 3.0 and SMRTbell Barcoded adapter 3.0 (Pacific Biosciences, Menlo Park, CA, USA). For each sample, DNA was sheared using g-tubes (Covaris, England) generating DNA fragments of approximately 9 kb. Size distribution was determined using a fragment Analyzer (Agilent Technologies, Santa Clara, CA, USA). Around 300 ng of each DNA sample were treated to remove single-strand overhangs and repair DNA damage. Ligation with overhang adapters to both double-stranded DNA ends was performed to create a closed, single-stranded circular DNA. After a nuclease treatment, size-selection was done on pooled samples to remove fragments less than 5Kb, with 3.1X 35% diluted AMPure PacBio beads. Quality and quantify of the libraries were determined using a Fragment Analyzer (Agilent Technologies) and a Qubit fluorimeter (Life Technologies). A ready-to-sequence SMRTBell Polymerase Complex was created using a Revio polymerase kit (PacBio). The PacBio Revio instrument was programmed to load a 300 pM library and sequenced in CCS mode on a PacBio SMRTcell 25M, with the Revio Sequencing Plate (Pacific Biosciences) and 24-h acquiring movie.

##### 1.2 Phylogenomic analysis of *Microcystis*

Normalised Robinson-Foulds metric [23] was used to assess the congruence between *Microcystis* phylogenetic tree and the dendrogram of its associated bacteria (AB). The phylogenetic tree was generated as described in the phylogenomic section of Methods (main manuscript). Associated bacteria presence/absence matrix was used to compute an AB jaccard distance matrix, using the vegdist function from the R library vegan (v2.7-1), for each taxonomic level (*i.e.* species, genus, family, order, class, phylum). Hierarchical clustering was performed on the AB distance matrix using the hclust function (R stats library v4.5.0), with complete linkage method, generating a dendrogram of the distance between *Microcystis* strains relatively to their associated bacteria. Robinson-Foulds' metric was then computed to compare topologies of the host phylogenetic tree and the AB dendrogram, using the R package Treedist (v2.11.0). Representation of the conserved splits was realized using the dendlist and tanglegram functions from the R package dendextend (v1.19.1).

For tanglegram representations involving the comparison of phylogeny to other hierarchical relationships, we pruned the phylogenetic tree, re-rooted it to get genospecies C as the outgroup, and made it ultrametric using phytools (v2.3_0) non-negative least square method.

#### 1.3 Functional annotation

We curated the identification of certain functions during metabolic network reconstruction of *Microcystis.* To handle potentially missing or mis-annotated functions, we systematically conducted a reciprocal-blast procedure. For the suspected false negatives, we blasted the entire MAGs on a local database of the researched functions [4]. We considered a protein sequence as homologous to one (preferably multiple) hits in the database if the following thresholds were respected: identity $\geq$ 80% and p-value $\leq$ 0.05. To double-check the protein characterisation, and avoid false positives, we then re-blasted it against the Uniprot database [6] and assessed whether another function was actually a better match. AntiSMASH outputs were a supplementary argument to support the identification of the putatively missing genes clusters.

#### 1.4 Genome-scale metabolic network comparison

GSMNs comparison was performed based on their reaction contents using PADMet-analysis [1]. To compare *Microcystis* GSMN to its phylogeny, a tanglegram was constructed on a hierarchical clustering using complete linkage method and Jaccard distance of reactions presence-absence matrix. To compare GSMN contents at the phycosphere level, we performed a principal coordinate analysis (PCoA, based on a Bray-Curtis dissimilarity) taking into account AB relative abundances. First, we computed reactions presence-absence matrix of all members per phycosphere, then weighted those presence using the genome relative abundance in the community according to metagenomic data. Finally, we summed the weighted values to obtain a value for each reaction in each phycosphere.

#### 1.5 Metabolic modelling

The modelling framework used by Metage2Metabo [3] relies on network expansion [9]. Briefly, each metabolite of the GSMN is either reached (producible) in the simulated conditions, or non-reached (unproducible). Network expansion states that products of a reaction are reached only if all of the reaction's reactants are reached or belong to the seeds. M2M distinguishes the metabolites that can be reached individually by GSMNs from those relying on metabolic complementarity and mutualistic interactions to be producible.

Metabolic modelling simulations require a list of compounds called *seeds* that initiate the reachability of molecules in the metabolic networks. Following the composition of BG11 growth medium used experimentally, Supp. Table **S1** describes the seeds used in the models.

Two types of simulations were performed. The first one is at the phycosphere level, simulating lab culture conditions of each community and estimating the *intra-phycosphere metabolic complementarity.* For each community, the respective GSMNs of the cyanobacterium and of its associated bacteria were gathered and M2M was run with the command `metacom`. The second simulation aimed at modelling the interactions among distinct phycospheres in lake-like natural conditions. For this, each collection of GSMNs related to a phycosphere was merged with the PADMet library, and the simulated community consisted of twelve phycosphere-level networks, enabling to measure the *inter-phycosphere metabolic complementarity* potential. Resulting metabolic potentials (reachable metabolites by the whole community or by individual community members) were then analysed. Producible compounds were grouped using the metabolite classification of natural products from [16].

| Name | Chemical Formula | MetaCyc ID |
|---|---|---|
| adp* | C10H12N5O10P2 | ADP |
| ammonium | NH4+ | AMMONIUM |
| biotin | C10H15N2O3S | BIOTIN |
| boric acid | H3O3B | BORATE |
| calcium | Ca+ | CA+2 |
| calcium chloride dihydrate | CaCl2 | CPD0-2507 |
| carbon dioxide | CO2 | CARBON-DIOXIDE |
| chloride | Cl | CL- |
| citrate | C6H5O7 | CIT |
| cobalt(II) | Co+2 | CO+2 |
| cobalt(II) nitrate | Co(NO3)2 | CPD-22943 |
| copper(II) | Cu+2 | CU+2 |
| copper(II) sulfate | Cu(SO4) | CUSO4 |
| coA* | C21H32N7O16P3S | CO-A |
| cyanocob(III)alamin | C63H88N14O14PCo | CPD-315 |
| dioxygen | O2 | OXYGEN-MOLECULE |
| dipotassium phosphate | K2HPO4 | CPD0-2433 |
| EDTA disodium salt | Na2(C10N2H14O8) | CPD-19640 |
| ferric ammonium citrate | Fe(NH4)(C6H5O7) | CPD-19639 |
| folate class* | | THF-GLU-N |
| hepes solution | C8H18N2O4S | HEPES |
| hydrogen chloride | HCl | HCL |
| iron(III) | Fe+3 | FE+3 |
| light | | Light |
| manganese(II) | Mn+2 | MN+2 |
| manganese(II) sulfate | Mn(SO4) | CPD0-1608 |
| molybdate | O4Mo | CPD-3 |
| nad* | C21H26N7O14P2 | NAD |
| nadp* | C21H25N7O17P3 | NADP |
| NAD ou NADP* | | NAD-P-OR-NOP |
| nitrate | NO3 | NITRATE |
| phosphate | PO4 | Pi |
| potassium | K+ | K+ |
| sodium | Na+ | NA+ |
| sodium molybdate dihydrate | Na2(MoO4)(H2O)2 | CPD-19634 |
| sodium nitrate | Na(NO3) | CPD-12921 |
| sulfate | SO4-2 | SULFATE |
| sulfide | H2S | HS |
| thiamine | C12H17N4OS | THIAMINE |
| thiamine HCl | (HCl)(C12N4H17SO) | CPD-22876 |
| ubiquinones* | | Ubiquinones |
| water | H2O | WATER |
| zinc | Zn+2 | ZN+2 |
| zinc sulfate | Zn(SO4) | CPD0-2393 |

**Supplementary Table S1**: **Adapted seed list from the BG11 culture medium**. An asterisk (∗) follows cofactors that are not part of BG11's composition, but are added *in silico* to initiate reachability, as they are common and important enough for basic metabolic process to be included in the seeds.

#### 1.6 Metabolomic data acquisition and analyses

##### 1.6.1 Monophasic (MP) and biphasic (BP) metabolomic extractions protocols

In the MP method, about 1 mg of lyophilised biomass or supernatant was extracted with 100 µL of solvent mix of methanol/water (3:1, v/v) acidified with 0.1 percent formic acid. Metabolite extraction was carried out on ice using an ultrasonic probe, performing four consecutive cycles of 30 seconds, each followed by a 30-second pause. The samples were then centrifuged (15,000 *g*, 4°C, 10 minutes). A volume of 50 µL of the supernatant was transferred into injection vials, and 1 µL was analysed by ultra-high-performance liquid chromatography (Elute, Bruker, Bremen, Germany) coupled with time of flight high-resolution tandem mass spectrometry (Maxis II-QTOF, Bruker, Bremen, Germany) (UHPLC-HRMS/MS).

In the BP method, about 2 mg of dried biomass was first extracted on ice by adding 1 mL of cold mixture of methyl-tert-butyl-ether and methanol (MTBE-MeOH, ratio 3:1, v:v) and homogenisation during 2 cycle of 15-second (6000 RPM, FastPrep-24 5G, MP-biomededical). Then, a 650 µL cold mixture of ultrapure water and methanol (UPW-MeOH, ratio 3:1, v:v) was added, followed by a third cycle of homogenization. Hydrophilic and lipophylic fractions were separated by centrifugation (12000 RPM, 4°C, 5 min) and further partly collected. A second similar extraction was performed to optimise metabolite recovery after the addition of 700 µL of MTBE-MeOH mixture and 455 $\mu L$ of UPW-MeOH mixture. In total, lipophilic and hydrophilic fractions consisted in 1.1 mL and 1.3 mL extracts, respectively. Following the extraction, 500 µL of each fractions were evaported to dryness (EZ2-PLUS, Genevac) and resuspended in ACN:H20 (1:1, v/v) mixture and ISO:ACN (1:1, v/v) respectively for hydrophilic and lipophilic fractions. Both fractions were analysed on UHPLC Vanquish combined to Q-Exactive Plus (Qex+) HRMS equiped with a heated electropray ionisation probe (HESI-II) (ThermoFisher Scientific, San Jose, CA).

##### 1.6.2 Elution gradients

MP extract was separated on a C18 Acclaim PolarAdvantage II column (2.1x100 mm, 2.2 µm, ThermoScientific), as described in Supp. Table **S2**.

| Time (min) | Flow (mL min$^{-1}$) | Eluent A (%) | Eluent B (%) |
|---|---|---|---|
| 0 | 0.300 | 95 | 5 |
| 2 | 0.3 | 95 | 5 |
| 16 | 0.300 | 10 | 90 |
| 18 | 0.300 | 10 | 90 |
| 19 | 0.300 | 95 | 5 |
| 21 | 0.300 | 95 | 5 |

**Supplementary Table S2**: **Gradient used to elute the monophasic extracts on C18 Acclaim PolarAdvantage II column at 40°C**. Eluent A, UPW supplemented with 0.08% of FA; Eluent B, ACN supplemented with 0.08% of FA.

BP extracts were separated on HSST3 RP-C18 column (150 × 2.1 mm × 1.7 µm, Waters) to analyse hydrophilic and lipophilic fractions (Supp. Tables **S3**, **S4**)) and BEH-Amide (100 x 2.1 mm x 1.7 µm, Waters) to analyse only hydrophilic fraction (Supp. Table **S5**).

| Time (min) | Flow (mL min$^{-1}$) | Eluent A (%) | Eluent B (%) |
|---|---|---|---|
| 0 | 0.400 | 99 | 1 |
| 4 | 0.400 | 99 | 1 |
| 20 | 0.400 | 1 | 99 |
| 22 | 0.400 | 1 | 99 |
| 22.1 | 0.400 | 99 | 1 |
| 25 | 0.400 | 99 | 1 |

**Supplementary Table S3**: **Gradient used to elute the hydrophilic fraction on HSST3 column at 45°C**. Eluent A, UPW supplemented with 0.1 % of FA ; Eluent B, ACN supplemented with 0.1 % of FA.

| Time (min) | Flow (mL min-1) | Eluent A (%) | Eluent B (%) |
|---|---|---|---|
| 0 | 0.300 | 70 | 30 |
| 4 | 0.300 | 55 | 45 |
| 22 | 0.300 | 30 | 70 |
| 24 | 0.300 | 1 | 99 |
| 26 | 0.300 | 1 | 99 |
| 30 | 0.300 | 70 | 30 |

**Supplementary Table S4**: **Gradient used to elute the lipophilic fraction on HSST3 column at 45°C**. Eluent A, UPW/ACN (60:40, v:v) supplemented with 0.1 % of FA ; Eluent B, ACN/isopropanol (10:90, v:v) supplemented with 0.1 % of FA.

| Time (min) | Flow (mL min$^{-1}$) | Eluent A (%) | Eluent B (%) |
|---|---|---|---|
| 0 | 0.35 | 20 | 80 |
| 2 | 0.35 | 20 | 80 |
| 12 | 0.35 | 80 | 20 |
| 16 | 0.35 | 80 | 20 |
| 18 | 0.35 | 20 | 80 |
| 24 | 0.35 | 20 | 80 |

**Supplementary Table S5**: **Gradient used to elute the hydrophilic fraction on BEH-Amide**. Eluent A, UPW supplemented with ammonium acetate 0.8% / EtOH (95:5; v:v) ; Eluent B, ACN supplement with ammonium acetate 0.8% / EtOH (95:5; v:v).

##### 1.6.3 Data acquisition

For MP, each extract was analysed in triplicate in positive ionization mode with a mass range of 50–1500 *m/z*, and automatic fragmentation of the TOP 5 most intense ions (autoMS/MS). QC and Blank samples (injected every six samples and every triplicate, respectively) were examined to ensure the reproducibility and robustness of the whole data series.

For BP, samples were also analyses in triplicates on QEx+ in both positive and negative mode according the parameters presented in Supp. Table **S6**.

| | DDA full scan (top 5) | **DDA MS/MS** |
|---|---|---|
| **Scan Range (m/z)** | 50-1500 | |
| **Resolution (FWHM)** | 70,000 | 17,500 |
| **Automated Gain Control (AGC)** | 3e6 | 1e5 |
| **Spray Voltage (kV)** | Pos : +4 ; Neg : -3.5 | 4 |
| **Sheath Gas Flow rate (µA)** | Pos : 40 ; Neg : 30 | 40 |
| **Capillary Temperature (°C)** | Pos : 350 ; Neg : 300 | 350 |
| **NCE** | - | 50 |
| **Loop Count** | - | 2 |

**Supplementary Table S6**: **Qex+ parameters for the analysis of BP extracts** after hydrophilic, lipophilic and HILIC separation

.

##### 1.6.4 Data processing, annotation and chemometrics

In order to highlight specialised metabolites, MetaboScape software(R) (Bruker Daltonics, Bremen, Germany) was first used to process only MP data. After recalibration of each sample analysis (< 1ppm, according to sodium formate internal standard), peaks with intensities greater than 5,000 counts in at least 10% of the set of samples and minimal RT-correlation coefficient of 0.7. Different states of charge (1+, 2+ and 3+ ) and classical adducts were grouped together, and the peak area was determined in order to generate a unique global data matrix containing semi-quantification results for each metabolite in all analysed samples. Feature annotations were attempted according to respective ion mass (< 2ppm) and isotopic pattern (< 20msigma) by automatic match with the CyanometDB 1.0 database [14] which contains over 2,100 chemical formulas of known specialised metabolite produced by cyanobacteria.

For untargeted metabolomics processing, MSDial software was used to extract features from samples. MSDial parameters for both MP and BP extracts are provided in Supp. Table **S7**.

| | **QTOF (Monophasic)** | **Qexactive+_ Orbitrap (Biphasic)** |
|---|---|---|
| MSDIAL version | v5.5.251021 | v5.5.251021 |
| **Conversion** | | |
| Software or package (specify language) | no | ThermoParser |
| Spectrum | - | mzML |
| MS levels | - | All |
| Peak Picking | - | All |
| Metadata | - | TXT |
| All detectors | - | Yes |
| Exceptions | - | Exclude reference and exception data |
| Gzipped | - | No |
| Compression | - | None |
| Errors | - | Ignore missing instrument properties |
| **Data Collection** | | |
| Mass accuracy | | |
| MS1 tolerance | 0.01 | 0.01 |
| MS2 tolerance | 0.05 | 0.05 |
| *Advanced - Data collection* | | |
| Rt time begin | 0 | 0 |
| Rt time range end | 100 | 100 |
| MS1 mass range begin | 0 | 0 |
| MS1 mass range end | 2000 | 2000 |
| MS/MS mass range begin | 0 | 0 |
| MS/MS mass range end | 2000 | 2000 |
| Excute Rt correction | NO | NO |
| *Advanced - Isotope recognition* | | |
| maximum number of isotopes | 5 | 5 |
| maximum number of charges | 1 | 1 |
| Consider Cl and Br Elements | YES | YES |
| *Multithreading* | | |
| Number of threads | 40 | 40 |
| **Peak Detection** | | |
| *Peak detection parameters* | | |
| Minimum peak height | 2.E+03 | 5.E+05 |
| Mass slice width | 0.1 | 0.1 |
| *Advanced* | | |
| Smoothing method | Linear weighet moving average | Linear weighet moving average |
| Smoothing level | 3 | 3 |
| Minimum peak width | 5 | 5 |
| Exclusion mass list | NO | NO |
| **Spectrum deconvolution** | | |
| *Deconvolution parameters* | | |
| Sigma window value | 0.5 | 0.5 |
| MS/MS abundance cut off | 25 | 10 |
| *Advanced* | | |
| Exlucde after precursor ion | YES | YES |
| Keep the isotopic ion until | 5 | 5 |
| Kepp the isotopic ions w/o MS2Dec | YES | YES |
| **Identification** | | |
| *Database setting* | | |
| Name | FragHub | FragHub |
| Last updated | 24-Apr-24 | 24-Apr-24 |
| Downloaded from | \href{https://zenodo.org/records/11057687}{https://zenodo.org/records/11057687 } | \href{https://zenodo.org/records/11057687}{https://zenodo.org/records/11057687 } |
| Comments | MS/MS positive mode | MS/MS positive mode |
| Comments | Combination of currated GNPS, Mslib, MassBank, Riken, Mona | Combination of currated GNPS, MsLib, MassBank, Riken, Mona |
| *Annotation Method* | | |
| Accurate Mass Tolerance (MS1) | 0.015 Da | 0.01 Da |
| Accurate Mass Tolerance (MS2) | 0.03 Da | 0.05 Da |
| Retention time Tolerance | 100 | 100 |
| *MS2 spectrum cut off* | | |
| spectrum amplitude cut off (relative) | 0 | 0 |
| spectrum amplitude cut off (absolute) | 0 | 0 |
| Mass range begin (Da) | 0 | 0 |
| Mass range begin (Da) | 2000 | 2000 |
| *Annotation cut off* | | |
| Dot product score | 600 | 600 |
| weighted dot product score | 600 | 600 |
| reverse dot product score | 800 | 800 |
| Matched spectrum percentage (%) | 50% | 25% |
| Minimum number of matched spectrum | 5 | 3 |
| *Retention time scoring* | | |
| Use retention time or scoring | no | no |
| use retention time for filtering | no | no |
| Adduct ion | | |
| Data setting | [M+H]+, [M+NH4]+, [M+Na]+, [M+CH3OH+H]+, [M+K]+, [M+Li]+, [M+ACN+H]+, [M+H-H2O]+, [M+H-2H2O]+, [M+2Na-H]+, [M+ACN+Na]+, [M+2K-H]+, [M+2ACN+H]+, [M-C6H8O6+H]+, [2M+H]+, [2M+NH4]+, [2M+Na]+, [2M+3H2O+2H]+, [2M+K]+, [2M+ACN+H]+, [2M+ACN+Na]+, [M-H]-, [M-H2O-H]-, [M+Na-2H]-, [M+FA-H]-, [M-C6H10O4-H]-, [M-C6H10O5-H]-, [M-C6H8O6-H]-, [M+CH3COONa-H]-, [2M-H]-, [2M+FA-H]-. | [M+H]+, [M+NH4]+, [M+Na]+, [M+CH3OH+H]+, [M+K]+, [M+Li]+, [M+ACN+H]+, [M+H-H2O]+, [M+H-2H2O]+, [M+2Na-H]+, [M+ACN+Na]+, [M+2K-H]+, [M+2ACN+H]+, [M-C6H8O6+H]+, [2M+H]+, [2M+NH4]+, [2M+Na]+, [2M+3H2O+2H]+, [2M+K]+, [2M+ACN+H]+, [2M+ACN+Na]+, M-H]-, [M-H2O-H]-, [M+Na-2H]-, [M+FA-H]-, [M-C6H10O4-H]-, [M-C6H10O5-H]-, [M-C6H8O6-H]-, [M+CH3COONa-H]-, [2M-H]-, [2M+FA-H]-. |
| **Alignment parameters** | | |
| *Alignment parameters setting* | | |
| Result Name | x | x |
| Reference File | First QC | First QC |
| Retention time tolerance | 0.2 | 0.2 |
| MS1 tolerance | 0.025 | 0.025 |
| *Advanced* | | |
| Retention time factor | 0.5 | 0.5 |
| MS1 factor | 0.5 | 0.5 |
| Peak count filter | 0 | 0 |
| N% detected in at least one group | 0 | 0 |
| remove features based on blank information | NO | NO |
| Sample max / blank average | 5 | 5 |
| Keep reference match metabolites features | YES | YES |
| Keep suggested (w/o MS2) metabolites features | YES | YES |
| Keep removable features and assigne the tag | YES | YES |
| gap filling by compulsion | YES | NO |
| *MScleanR* | | |
| Blank ratio | 0.8 | 0.8 |
| Delete Ghost Peak | YES | YES |
| Incorrect Mass | YES | YES |
| Relative Standard Deviation (max RSD) | 50 | 50 |
| Relative Mass Defect | 50-3500 | 50-3500 |

**Supplementary Table S7**: **List of parameters in MS-DIAL** (v5.25) for processing metabolomics raw data and further filtering

Feature intensity was further normalised by the Total Ion Chromatogram (TIC) prior the use of MS-cleanR [10] to filter the data according blank intensity, RSD, RMD, etc. (parameters in Supp. Table **S7**.) Filtered data were finally export as peak lists (i.e. one per acquisition mode) for chemometrics analysis and as spectra (as individual .mat files) to implement SIRIUS (v6.1) [7]. SIRIUS software allowed to interrogate structural libraries (SIRIUS database and cyanoMetDB [15]) and provide scoring of the formula (ZODIAC) and structure (CSI:FingerID). [8, 17, 26]. In has to be noted, that these filtered peak lists contained both "only MS1" features (i.e. detected signals for which no fragmentation was performed) and "MS2" features (i.e. signals for which a fragmentation was performed allowing to interrogate spectral or structural DBs).

Output files from MS-DIAL and SIRIUS (Top 50 per feature) were further imported into MS-Net (developed in Knime Analytics Platform, v5.2) [11] in order to merge Pos and Neg acquisition for each gradient (hydrophilic-HSST3-pos/neg, hydrophilic-HILIC-pos/neg, lipophilic-pos/neg), suppress analytical redundancy, provide annotation confidence level and classification (NPclassifier, [16]), and highlight link between metabolites and identified bacterial genus and *Microcystis* sp. Briefly, among the top 50 in silico candidates per feature, those matching taxonomic criteria (genus: *Gemmatimonas, Hydrogenophaga, Meiothermus, Neoroseomonas, Aminobacter, Limnobacter, Novosphingobium, Phenylobacterium, Nevskia, Rhabdaerophilum, Cnuella, Flavobacterium, Aquidulcibacter, Brevundimonas, Caulobacter, Silanimonas, Inhella, IdeonellaA, Pararheinheimera, Falsiroseomonas, PseudomonasE, Rubrivivax, LeptospiraA, Microcystis* ; Famillies : *Fimbriimonadaceae, Sphingomonadaceae, Gemmatimonadaceae, TH1-2, Burkholderiaceae, Thermaceae, Acetobacteraceae, Caulobacteraceae, Rhizobiaceae, Nevskiaceae, Beijerinckiaceae, Chitinophagaceae, Flavobacteriaceae, Tepidisphaeraceae, Xanthomonadaceae, Alteromonadaceae, Pseudomonadaceae, Usitatibacteraceae, Leptospiraceae, Microcystaceae*) were elevated to Level 3a.

Features were filtered using Ion Identity Network (IIN) results to remove redundant adducts and isotopes, followed by MS-CleanR-based RT clustering ($\Delta$RT $\leq$ 0.01 min). Within each cluster, the top 2 features by network degree and the top 2 by peak intensity were retained. The MS2 network was constrained to edges with cosine similarity $\geq$ 0.7 and $\Delta$RT $\leq$ 8 min between connected nodes. High-confidence annotations (Levels 1, 2a, 3a) seeded the MS2 network for iterative annotation propagation. For each feature pair, candidate structures were ranked using a weighting parameter set to $\alpha = 0.3$, prioritizing structural-spectral evidence (70%) over *in silico* ranking (30%). The top 5 candidates by Link Score were retained per feature before looping through the entire MS2 network. Redundant annotations with identical InChIKey identifiers and Pearson correlation among samples $>$ 0.7g were consolidated by selecting the candidate with the highest mean peak height. Positive and negative mode feature lists were merged using $\Delta$RT $\leq$ 0.05 min, $\Delta$m/z $\leq$ 0.005 Da, and a minimum Pearson correlation $\geq$ 0.6 across sample intensities. Final annotations were enriched with chemical ontology classifications from ClassyFire (kingdom, superclass, class, subclass) and NPClassifier (pathway, superclass, class). Database identifiers (PubChem CID, KEGG, HMDB, ChEBI) were retrieved using the Chemical Translation Service. Natural product-likeness scores were calculated using the NPlikeness calculator [13]. Structural similarity networks were constructed using Tanimoto coefficient $\geq$ 0.8, retaining the top 2 nearest neighbours per feature.

Prior chemometrics, replicates were combined through averaging as to keep one signal per metabolite for each phycosphere. Then, all the peak lists (MP, BP-hydro-HSST3-pos/neg, BP-hydro-HILIC-pos/neg, BP-lipo-pos/neg) were merged and filtered in order to remove redundancy by keeping single annotation based on InChIKey, while features with no MS2 might present redundancy (i.e. adduct from the same molecules). Feature intensity was normalised, transformed and scaled (*i.e.* Sum normalisation, cube root transformation and pareto scaling) before statistical analyses.

#### 1.7 Comparison of GSMN contents and metabolic models with metabolomics

Mapping of the entire metabolomic dataset to GSMN contents (including the PGs) was performed with MetaNetMap (v1.0.1) [20] using MetaCyc (v.29.0) as a reference mapping database. Metabolite presence-absence data per phycosphere was obtained by intersecting metabolomic signal (requiring detection in $\geq$ 2 out of 3 replicates per community) with GSMN mapping. Network hit proportions were calculated as the number of networks mapped per metabolite divided by the number of members in the phycosphere. A heatmap was generated with communities as rows and metabolites as columns, stratified by NP-Classifier classes.

In addition, Venn diagram was generated using Jvenn [2] in order to compare annotated metabolites from metabolomics and predicted metabolites from GSMNs based on InChiKey.

Finally, the chemical landscape of *Phyco1364* was summarised as sunplot according the number of features per NPC_pathway/superclasses/classes and further compared to this predicted from GSMNs modelling.

### 2 Supplementary results

#### 2.1 Taxonomy of *Microcystis*' associated microbiome

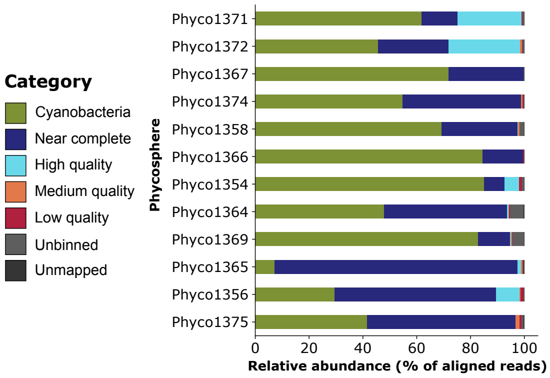


**Supplementary Figure S1**: **Relative abundance of Cyanobacteria, ABs (near complete, high, medium, or low quality MAGs), or unbinned contigs** according to metagenomic read mapping.

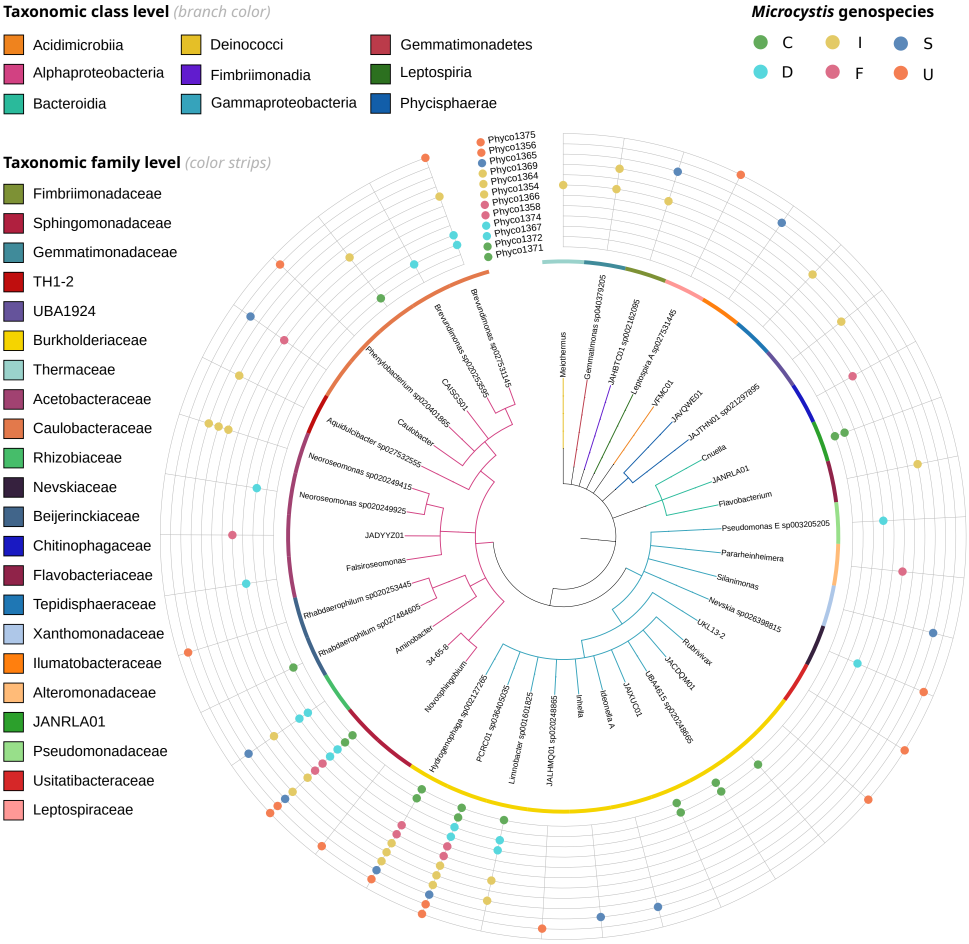


**Supplementary Figure S2**: **Phycospheres bacteria composition.** Taxonomic tree covering all distinct taxa identified from phycospheres high quality MAGs. The tree outer coloured circle annotates the Family of the bacteria, the branches are coloured by taxonomic Classes. On the very outer part, the grid identifies which taxa are found in which phycosphere; a coloured marble corresponding to the phycosphere is present when the taxa was annotated at least once in it.

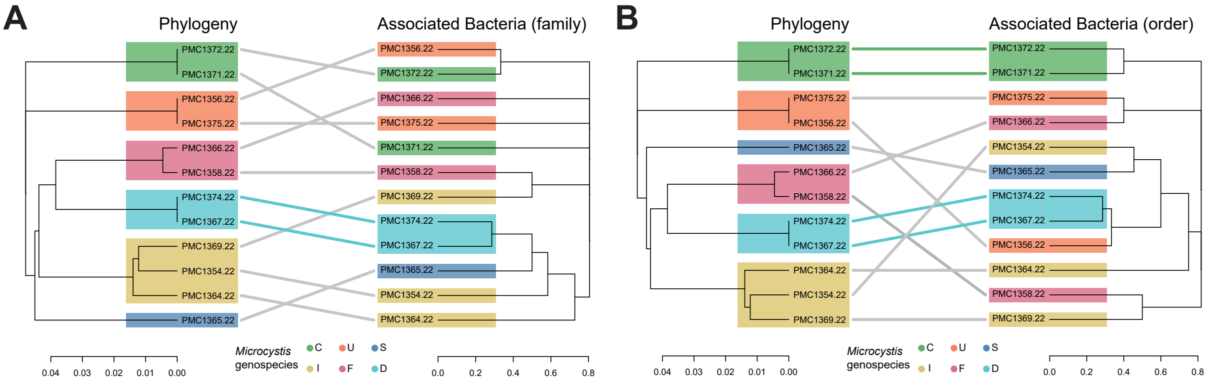


**Supplementary Figure S3**: **Relationships between *Microcystis* strain phylogenomy and the taxonomic composition of the respective associated bacteria according to hierarchical classification A.** Family taxonomic level. **B.** Order taxonomic level.

#### 2.2 Microcystin-LR biosynthesis pathway

| Enzyme | Type | Main function | References | *Microcystis* with this gene |
|---|---|---|---|---|
| McyG | PKS-NRPS | Activation of initiator phenylpropanoid; elongation by Malonyl-CoA | [22, 12, 27] | 5 |
| McyJ | O-MT | O-methylation | [5, 27] | 5 |
| McyD | PKS | Elongation by Malonyl-CoA; Dehydration | [27] | 5 |
| McyE | PKS-NRPS | Elongation by D-Glu | [27] | 5 |
| McyF | Glutamate Racemase | Epimerization of L-Glu into D-Glu | [27, 24] | 7 |
| McyA | NRPS | Elongation by L-Ser and D-Ala; Epimerization of L-Ala into D-Ala | [27] | 5 |
| McyI | 2-HADH | Production of D-MeAsp | [19, 21] | 5 |
| McyB | NRPS | Elongation by L-Leu and D-MeAsp | [27] | 5 |
| McyC | NRPS | Elongation by L-Arg; Cyclization | [27] | 5 |
| McyH | ABC transporter | Extracellular transport of MC-LR | [27] | 5 |

**Supplementary Table S8**: **Description of the enzymes involved in the biosynthesis of microcystin-LR**, in their order of action. **Abbreviations**: NRPS, Non-Ribosomal Peptide Synthetase; PKS, Polyketide Synthase; ABC transporter, ATP-Binding Cassette transporter; HADH, Hydroxy Acid Dehydrogenase; MT, O-methyltransferase.

#### 2.3 Phycosphere metabolic diversity

##### 2.3.1 Metabolic annotations of associated bacteria (AB)

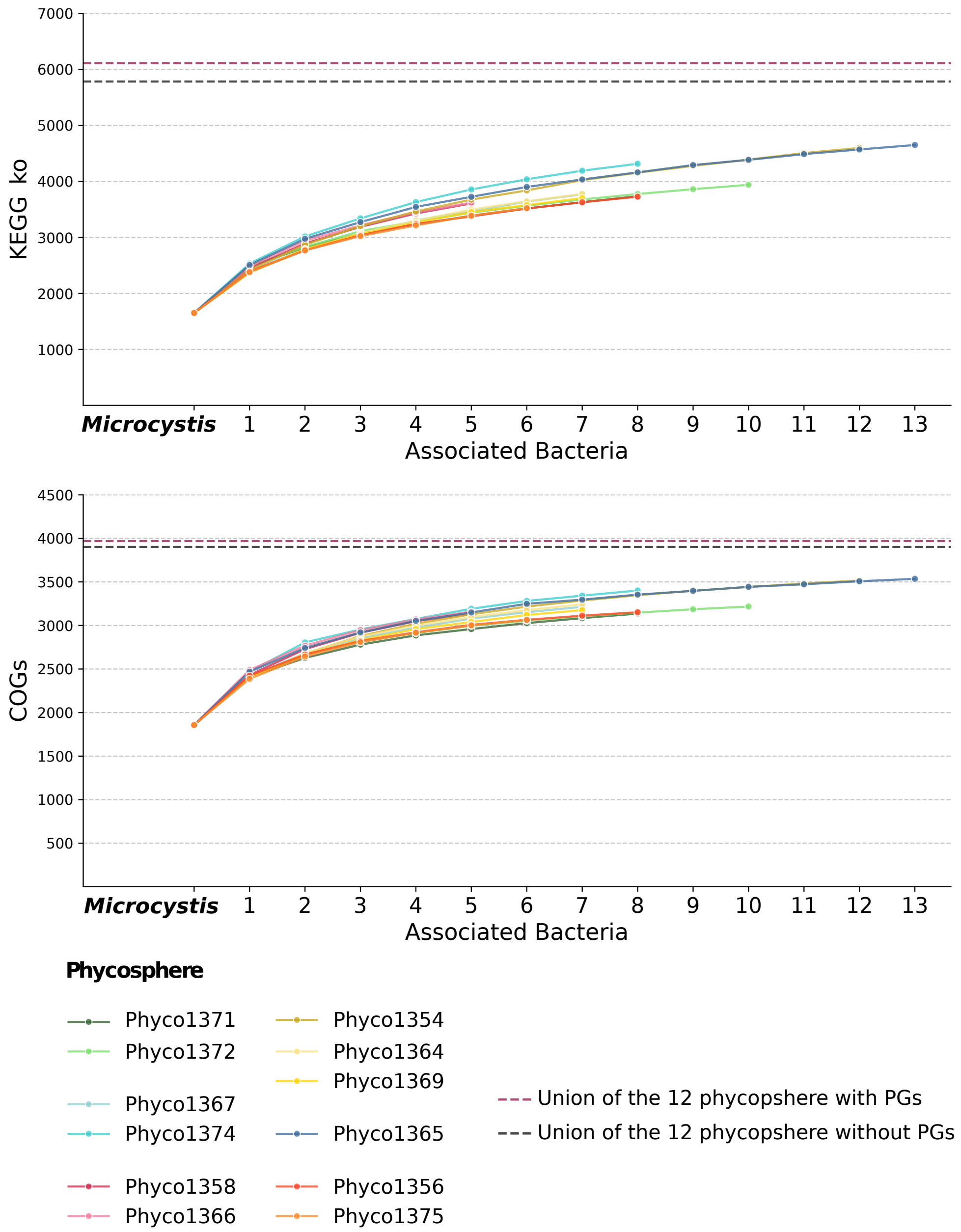


**Supplementary Figure S4**: **Rarefaction-curve of KEGG KO (above) and COGs (below) annotations** from phycosphere metagenomes. Rarefaction curves represent accumulation of unique annotations as a function of the number of sampled associated bacteria, based on 100 sampling iterations. The starting point is *Microcystis*. Pseudogenomes (PGs) are excluded from the sampling of phycosphere's individual curves.

##### 2.3.2 Biogechemical cycles

**Nitrogen Cycle** For almost all phycospheres, at least one associated bacterium—and not *Microcystis*—carried the nitrate reductase genes *narGH* (EC:1.7.5.1), as well as nitrite reductase genes *NirBD* (EC:1.7.1.15) in every phycosphere. However, the nitrite reduction gene *NirA* (EC:1.7.7.1) was found in all *Microcystis* genomes. The NO-forming nitrite reductase gene *nirK* was detected in members of *Phyco1354*, *Phyco1365*, *Phyco1367*, *Phyco1374*, and *Phyco1375*, but not in *Microcystis*. Notably, these same phycosphere-associated bacteria were also the only ones to possess nitrite oxide reductase genes *NorBD* (EC:1.7.2.5). No genes for nitrogen fixation were identified in any MAGs, either from *Microcystis* or the heterotrophic bacteria.

**Methanogenesis** We did not detect *McrABC* genes for methanogenesis in *Microcystis* genomes, although *Microcystis* are suspected to produce methane [29]. However, a gene for methane oxidation—the methane monooxygenase regulatory protein B—was found in non-cyanobacterial members in nine out of twelve phycospheres.

**Complex Carbon** As the bacteria associated with *Microcystis* in phycospheres are heterotrophic, they are expected to carry genes for the degradation of complex carbohydrates such as oligosaccharides. Indeed, in some phycospheres, genes for xylose, mannose, and galactose degradation were identified. Additionally, genes implicated in the breakdown of biomolecules originating from phytoplankton and fungi—including cellulose, hemicellulose, and pullulan—were also detected.

### 3 Metabolomic analyses

#### 3.1 Added value of combined extraction protocols

Figure **S5** depicts the annotated (InChiKey identifiers) metabolomic signals obtained with each extraction protocol, highlighting their complementarity to provide a comprehensive picture of the phycosphere endometabolome.

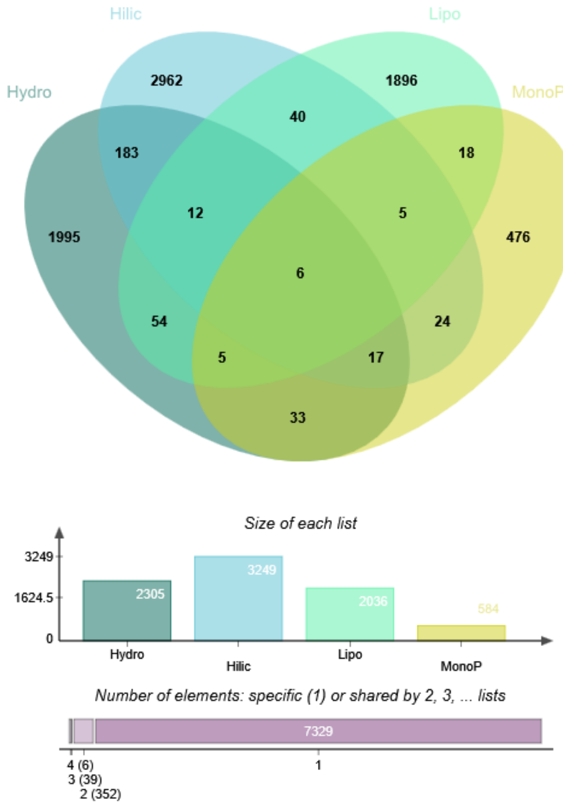


**Supplementary Figure S5**: **Venn diagram of putatively annotated features according to extraction protocols and chromatographic method**. The Venn diagram was generated with *jvenn* (`https://jvenn.toulouse.inra.fr/app/index.html`. **Abbreviations:** Hydro, Hydrophilic fraction separated on HSST3 Column; Hilic, Hydrophilic fraction separated on BEH-Amide column; Lipo, Lipophilic fraction separated on HSST3; MonoP, MP extract separated on C18 polar Advances II column.

#### 3.2 Biochemical composition of the phycospheres

The PCA in Supp. Fig. **S6** shows a separation of the phycosphere regarding their endometabolome composition only partly reflecting *Microcystis* genotype, as noted in the HCA (Main Figure 4A of the manuscript).

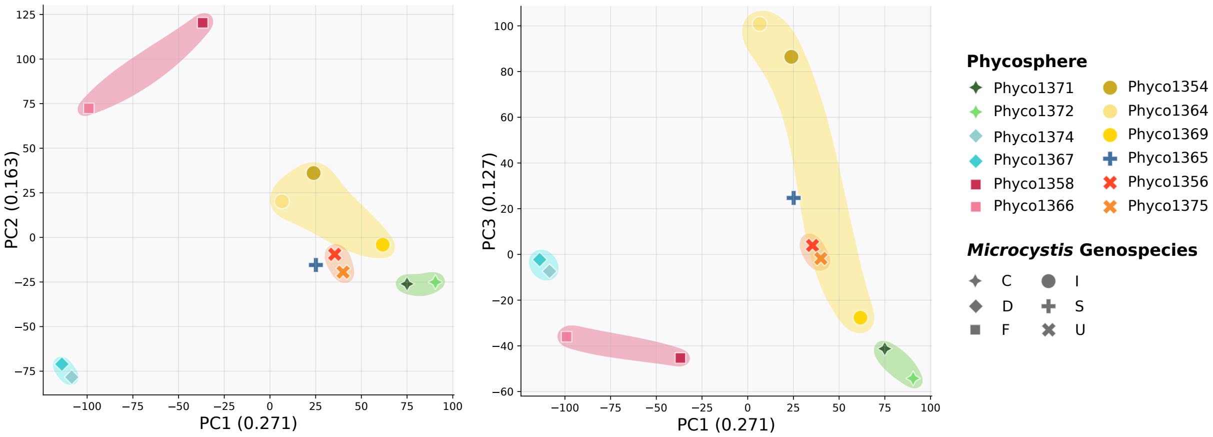


**Supplementary Figure S6**: **PCA of the transformed metabolomics signals.** For each of the extraction methods, triplicates were merged to get one signal per metabolite, and all extractions results aggregated. Coloured areas highlight phycosphere's associated to the same *Microcystis* genospecies.

Distribution of chemical classes (NPclassifier, [16]) relative intensity among the physcopheres are reported in Fig. **S7**. At the NPC-pathway level, the annotations do not showed marked differences in the biochemical composition of the twelve phycospheres.

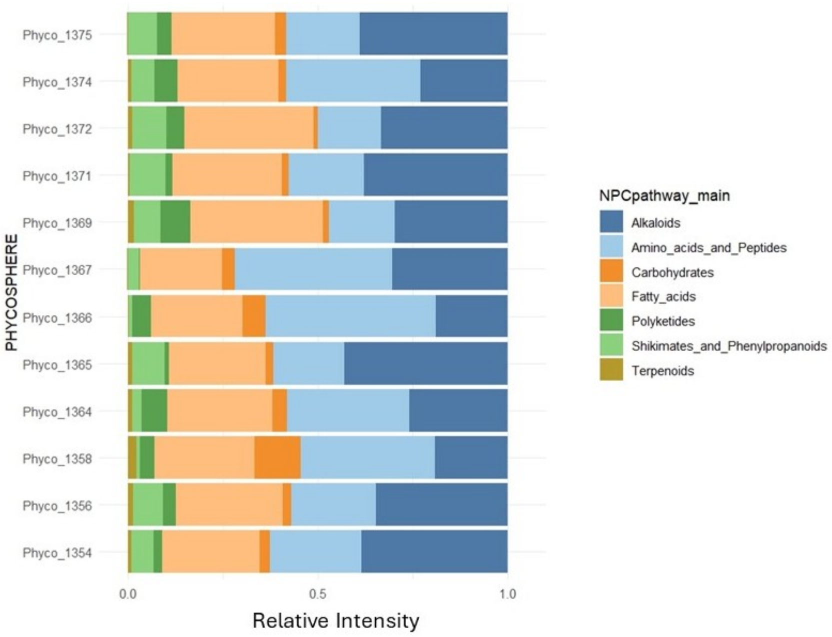


**Supplementary Figure S7**: **Barplot depicting the relative intensity of NPC Pathways among the phycospheres**.

#### 3.3 Taxonomic assignments of metabolomic signals

MS-Net highlights that among the 6016 putatively annotated and classified metabolites, 144 were reported in some taxa composing the phycosphere from metagenomics (Fig. **S8**). The main represented genus and familly was Microcystis and Microcystae (63 features), likely in accordance with the higher abundance of this taxa in most of the Phycosphere and also the higher number of studies on

Microcystins specialised metabolites in the literature, on which COCONUT database is based. Then, 40 features were reported with *Pseudomonas* or *Pseudomonoceae* and 10 with *Burkholderiaceae*. Among them, two apocarotenoids (C30) were putatively annotated. These compounds are known to have antioxidant properties [25]. We also annotated compounds with bactericidal effects like Sorbistin A2 [28] or siderophores like Ornibactin C6 and C8 [18]. These compounds may offer an ecological advantage in productive phycospheres. However, as they are annotated *in silico*, these results should be treated with caution.

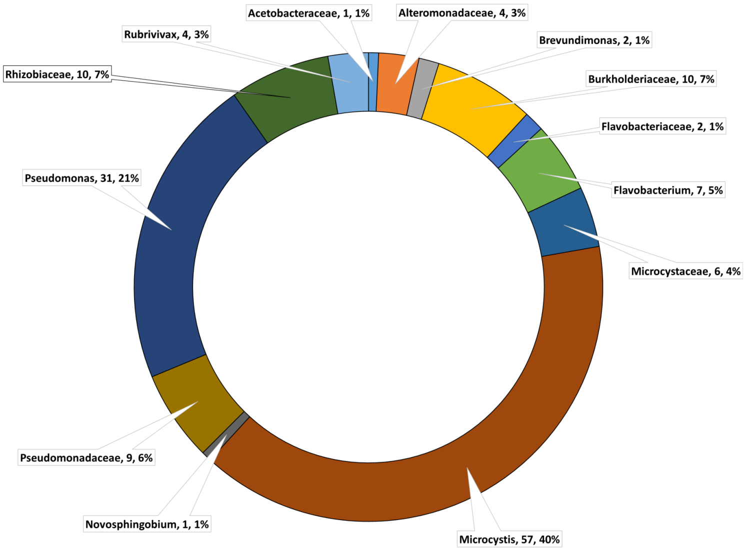


**Supplementary Figure S8**: **Circular plot depicting the number of features matching the Genera and the Families defined in MSnet parameters**. The percentage is from the total of 144 putatively annotated metabolites matching the taxa.

### 4 Metabolic modelling

#### Metabolic potential of associated bacteria and *Microcystis*

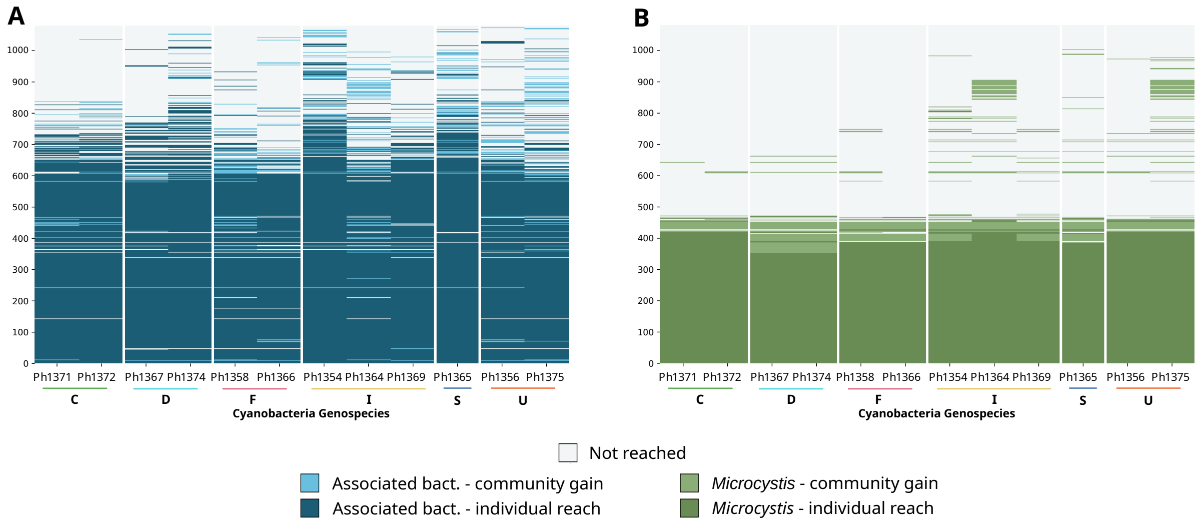


**Supplementary Figure S9**: **Modelling the metabolic complementarity intra-phycosphere A.** Heatmap of metabolites reached by the associated bacteria of each phycosphere grouped by genospecies (C, D, F, I, S, and U). **B.** Heatmap of metabolites reached by *Microcystis* in each phycosphere. "Individual reach" indicates that the metabolic capabilities of one member (either an associated bacterium in A, or the cyanobacterium in B) are sufficient to produce the metabolite from the growth medium. "Community gain" indicates that a member can reach the metabolites but only through metabolites exchanges, i.e. predicted cross-feeding interactions. "Not reached" depicts metabolites which are not predicted to be produced by the corresponding bacteria or cyanobacteria in the simulation, or are possibly absent from the corresponding GSMNs.

#### Comparison between intra-phycosphere model predictions and metabolomic signals

Comparison with merged metabolomics datasets revealed high gap between all predicted metabolites (1,078 metabolites with an InChiKey) and those actually annotated (7,696, L2-L3), as only 28 of the compounds predicted as producible could be clearly identified in the measurements. We then inspected steroids and hopanoids that were annotated through metabolomics, while their precursors, such as squalene, appeared absent. Conversely, metabolic modelling indicated that squalene can be produced in the community by certain *Microcystis* species. For the hopanoid bacteriohopanetetrol, a bacterial membrane compound annotated *in silico* in metabolomics, we identified its precursors (squalene and hopan-22(29)-ene) in the model predictions but not the final product. The presence of complete biosynthetic pathways in cyanobacterial GSMNs, along with precursors but not final products that are nevertheless detected by metabolomics suggests that modelling may preferentially reveal biosynthetic intermediates, thus partly explaining discrepancies between modelling predictions and metabolomics data.

[5] Guntram Christiansen, Jutta Fastner, Marcel Erhard, Thomas Börner, and Elke Dittmann. Microcystin biosynthesis in planktothrix: Genes, evolution, and manipulation. 185(2):564–572. Publisher: American Society for Microbiology.

[6] The UniProt Consortium, Alex Bateman, Maria-Jesus Martin, Sandra Orchard, Michele Magrane, Aduragbemi Adesina, Shadab Ahmad, Emily H Bowler-Barnett, Hema Bye-A-Jee, David Carpentier, Paul Denny, Jun Fan, Penelope Garmiri, Leonardo Jose da Costa Gonzales, Abdulrahman Hussein, Alexandr Ignatchenko, Giuseppe Insana, Rizwan Ishtiaq, Vishal Joshi, Dushyanth Jyothi, Swaathi Kandasaamy, Antonia Lock, Aurelien Luciani, Jie Luo, Yvonne Lussi, Juan Sebastian Martinez Marin, Pedro Raposo, Daniel L Rice, Rafael Santos, Elena Speretta, James Stephenson, Prabhat Totoo, Nidhi Tyagi, Nadya Urakova, Preethi Vasudev, Kate Warner, Supun Wijerathne, Conny Wing-Heng Yu, Rossana Zaru, Alan J Bridge, Lucila Aimo, Ghislaine Argoud-Puy, Andrea H Auchincloss, Kristian B Axelsen, Parit Bansal, Delphine Baratin, Teresa M Batista Neto, Marie-Claude Blatter, Jerven T Bolleman, Emmanuel Boutet, Lionel Breuza, Blanca Cabrera Gil, Cristina Casals-Casas, Kamal Chikh Echioukh, Elisabeth Coudert, Beatrice Cuche, Edouard de Castro, Anne Estreicher, Maria L Famiglietti, Marc Feuermann, Elisabeth Gasteiger, Pascale Gaudet, Sebastien Gehant, Vivienne Gerritsen, Arnaud Gos, Nadine Gruaz, Chantal Hulo, Nevila Hyka-Nouspikel, Florence Jungo, Arnaud Kerhornou, Philippe Le Mercier, Damien Lieberherr, Patrick Masson, Anne Morgat, Salvo Paesano, Ivo Pedruzzi, Sandrine Pilbout, Lucille Pourcel, Sylvain Poux, Monica Pozzato, Manuela Pruess, Nicole Redaschi, Catherine Rivoire, Christian J A Sigrist, Karin Sonesson, Shyamala Sundaram, Anastasia Sveshnikova, Cathy H Wu, Cecilia N Arighi, Chuming Chen, Yongxing Chen, Hongzhan Huang, Kati Laiho, Minna Lehvaslaiho, Peter McGarvey, Darren A Natale, Karen Ross, C R Vinayaka, Yuqi Wang, and Jian Zhang. UniProt: the universal protein knowledgebase in 2025. *Nucleic Acids Research*, 53(D1):D609–D617, 2024.

[7] Kai Dührkop, Markus Fleischauer, Marcus Ludwig, Alexander A Aksenov, Alexey V Melnik, Marvin Meusel, Pieter C Dorrestein, Juho Rousu, and Sebastian Böcker. Sirius 4: a rapid tool for turning tandem mass spectra into metabolite structure information. *Nature methods*, 16(4):299–302, 2019.

[8] Kai Dührkop, Huibin Shen, Marvin Meusel, Juho Rousu, and Sebastian Böcker. Searching molecular structure databases with tandem mass spectra using csi:fingerid. *Proceedings of the National Academy of Sciences*, 112(41):12580–12585, 2015.

[9] Oliver Ebenhöh, Thomas Handorf, and Reinhart Heinrich. Structural analysis of expanding metabolic networks. *Genome informatics. International Conference on Genome Informatics*, 15(1):35–45, 2004.

[10] Ophélie Fraisier-Vannier, Justine Chervin, Guillaume Cabanac, Virginie Puech, Sylvie Fournier, Virginie Durand, Aurélien Amiel, Olivier André, Omar Abdelaziz Benamar, Bernard Dumas, Hiroshi Tsugawa, and Guillaume Marti. MS-CleanR: A feature-filtering workflow for untargeted LC–MS based metabolomics. *Analytical Chemistry*, 92(14):9971–9981, 2020.

[11] Pereira Francisco, Duthen, Crossay, Alignan, Hennechart, Perez, and Marti. MS-net: Multi-similarity based network annotation for untargeted metabolomics. *Research Square*, 2025.

[12] Leslie M. Hicks, Michelle C. Moffitt, Laura L. Beer, Bradley S. Moore, and Neil L. Kelleher. Structural characterization of in vitro and in vivo intermediates on the loading module of microcystin synthetase. 1(2):93–102. Publisher: American Chemical Society.

[20] Coralie Muller, Juliette Audemard, Sylvain Prigent, and Clémence Frioux. Metanetmap: automatic mapping of metabolomic data onto metabolic networks. *bioRxiv*, 2026.

[21] Leanne A. Pearson, Kevin D. Barrow, and Brett A. Neilan. Characterization of the 2-hydroxy-acid dehydrogenase McyI, encoded within the microcystin biosynthesis gene cluster of Microcystis aeruginosa PCC7806. *The Journal of Biological Chemistry*, 282(7):4681–4692, February 2007.

[22] Kenneth L. Rinehart, Michio Namikoshi, and Byoung W. Choi. Structure and biosynthesis of toxins from blue-green algae (cyanobacteria). *Journal of Applied Phycology*, 6(2):159–176, April 1994.

[23] D. F. Robinson and L. R. Foulds. Comparison of phylogenetic trees. *Mathematical Biosciences*, 53(1):131–147, 1981.

[24] Heike Sielaff, Elke Dittmann, Nicole Tandeau De Marsac, Christiane Bouchier, Hans Von Döhren, Thomas Börner, and Torsten Schwecke. The mcyF gene of the microcystin biosynthetic gene cluster from microcystis aeruginosa encodes an aspartate racemase. 373:909–916.
